# A CRISPR-Cas9 screen Reveals STEEP1 as a Key Host Dependency Factor for Epstein-Barr Virus Latent Membrane Protein 1 Trafficking and Signaling

**DOI:** 10.64898/2026.03.19.712937

**Authors:** Shunji F. Li, Yizhe Sun, Yifei Liao, Ling Zhong, Eric M. Burton, Bidisha Mitra, Benjamin E. Gewurz

## Abstract

The Epstein-Barr virus (EBV) oncogene Latent membrane protein 1 (LMP1) is essential for B-cell transformation into continuously growing lymphoblastoid cell lines. LMP1 traffics to plasma membrane and intracellular signaling sites to mimic aspects of signaling by the B cell co-receptor CD40. LMP1 is expressed in many EBV-associated cancers, including post-transplant lymphoma, Hodgkin lymphoma, T/NK lymphoma and nasopharyngeal carcinoma, where it activates key growth and survival pathways. LMP1 signaling is also implicated in multiple sclerosis pathogenesis. To identify host dependency factors that support LMP1 trafficking and signaling, we performed a human genome-wide CRISPR-Cas9 screen in B cells. The screen identified both known and previously uncharacterized mediators of LMP1 signaling. The ER resident protein STEEP1, implicated in DNA sensor STING trafficking and signaling, was a top screen hit. Importantly, STEEP1 did not score in our prior B cell CRISPR screen for factors that support CD40 signaling, suggesting specificity. STEEP1 depletion strongly impaired LMP1 signaling, including activation of NF-kB and MAP kinase pathways.

Mechanistically, STEEP1 associated with LMP1 in a manner dependent on the N-terminal cytoplasmic tail and supported LMP1 egress from the ER to signaling sites in both B and epithelial cells. Collectively, these findings reveal STEEP1 as a key host factor that supports trafficking of newly synthesized LMP1 molecules to intracellular signaling sites and highlights LMP1/STEEP1 interaction as a novel therapeutic target.

**Importance:** Epstein-Barr virus (EBV) infects most people worldwide. While infection is often benign, it causes infectious mononucleosis, is associated with a range of lymphomas, nasopharyngeal and gastric carcinoma and is a major trigger for autoimmune disease, including multiple sclerosis. The EBV encoded oncogene LMP1 is a key driver of EBV pathogenesis, and its signaling is necessary for viral immortalization of B lymphocytes into continuously growing lymphoblasts (LCLs). Here, we performed a CRISPR genetic screen to identify host factors that support continuous, ligand-independent signaling by LMP1. This analysis identified an ER-resident protein called STEEP1, previously implicated in support of trafficking of the DNA sensor STING, as a key LMP1 partner.

We found that STEEP1 associates with LMP1 and supports LMP1 trafficking out of the endoplasmic reticulum to cellular signaling sites. As STEEP1 knockout impaired LMP1 function and LCL survival, our study identifies the STEEP1/LMP1 complex as a therapeutic target.

## Introduction

The gamma-herpesvirus Epstein-Barr virus (EBV) establishes lifelong infection in 95% adults worldwide [1, 2]. While infection is often benign, EBV is highly oncogenic and is causally linked to a diverse spectrum of malignancies, including multiple types of lymphomas, nasopharyngeal carcinoma, and gastric cancers [3–7]. Latent Membrane Protein 1 (LMP1) is frequently expressed in post-transplant lymphoproliferative disease (PTLD), central nervous system lymphoma, Hodgkin lymphoma, in T and NK lymphomas, in a subset of diffuse large B-cell lymphoma, in T/NK lymphomas and in nasopharyngeal carcinoma [8, 9]. EBV is also a major trigger of autoimmunity, in particular of multiple sclerosis [10] and systemic lupus erythematosus [11–13]. LMP1 expression may also be an important driver of autoimmunity, in particular with multiple sclerosis [14, 15].

LMP1 is a key EBV-encoded oncogene that mimics aspects of signaling by the B cell co-receptor CD40 [9, 16, 17]. LMP1 is essential for EBV-mediated transformation of primary B lymphocytes into continuously proliferating lymphoblastoid cell lines (LCLs), a key model for PTLD [18–20]. LMP1 expression is sufficient to induce oncogenic transformation of rodent fibroblast cell lines [21–23] and also drives B-cell proliferation, in particular when co-expressed with LMP2A and upon T and NK-cell depletion [9].

LMP1 constitutively signals in a ligand-independent manner [8, 24, 25]. LMP1 is comprised of a 24-residue N-terminal cytoplasmic tail, six transmembrane (TM) domains and a 200-residue C-terminal cytoplasmic tail. The TM domains drive LMP1 oligomerization and ligand-independent signaling from the C-terminal tail [26–28] [16, 17, 20, 29–33]. LMP1 activates multiple downstream pathways, including NF-κB, MAP kinase, interferon regulatory pathway and PI3 kinase pathways [19, 34–41]. LMP1 signaling from multiple C-terminal tail domains is required for LCL establishment [18, 42–44], and for LCL growth and survival [19].

LMP1 signaling is largely dependent on LMP1 trafficking out of the endoplasmic reticulum (ER) to plasma membrane and intracellular signaling sites, in particular lipid rafts [18, 26, 31, 45–48]. Knowledge remains incomplete about host factors important for LMP1 trafficking, including from the ER to Golgi. LMP1 targets a wide range of host genes, including those involved in cell survival [8, 19, 34, 49–52].

Here, we performed a human genome-wide CRISPR-Cas9 screen for host factors important for LMP1 signaling in Burkitt lymphoma B cells. These included well-characterized mediators of LMP1 NF-kB pathway activation as well as factors not previously implicated in LMP1 function. We provide multiple lines of evidence that the host ER-resident protein STEEP1 associates with LMP1, is a key dependency factor for LMP1 trafficking out of the ER and that therefore supports LMP1 signaling.

## Results

### Human genome-wide CRISPR-Cas9 screen for host factors that support LMP1 signaling

To identify host factors important for LMP1 function, we generated Cas9+ Daudi Burkitt B cells with conditional doxycycline (Dox)-inducible LMP1 expression, as recently described [19]. We used Daudi for this study as LMP1 signals robustly in them, but they are not dependent on LMP1 for survival, permitting genetic analysis of host factors important for LMP1 function. We recently characterized LMP1 host gene targets in Daudi cells[19], which have relatively low basal NF-κB activation, permitting detection of LMP1 effects on LMP1/NF-kB pathway targets. By contrast, LCLs are highly dependent on LMP1 for survival and rapidly undergo apoptosis upon loss of LMP1 signaling[19]. As a control, we also engineered cells to express doxycycline-inducible green fluorescence protein (GFP), such that doxycycline addition rapidly induced expression of both LMP1 and GFP. Furthermore, we previously used Daudi cells for a human genome-wide CRISPR screen for B cell factors important for CD40 signaling, in which cells were activated by addition of CD40-ligand [53, 54], enabling direct cross-comparison.

To screen for host factors important for LMP1 function, we transduced Cas9+ Daudi cells with the Brunello single guide RNA (sgRNA) library [55]. Brunello is comprised of 76,441 lentiviruses, each of which expresses one sgRNA. Across the library, human genes are targeted on average by four independent Brunello sgRNAs. To minimize transduction by multiple lentiviruses, 130 million Daudi cells were infected at a multiplicity of infection of 0.3. Five days post-transduction, B cells were treated with 200 ng/mL Dox for 16 hours to induce LMP1 and control GFP expression. As a readout of LMP1 signaling, we stained cells for FAS, whose expression is low on Daudi cell plasma membranes, but which is rapidly induced by LMP1 signaling [19, 52, 56–59] (**Fig. 1A**).

**Figure 1.**
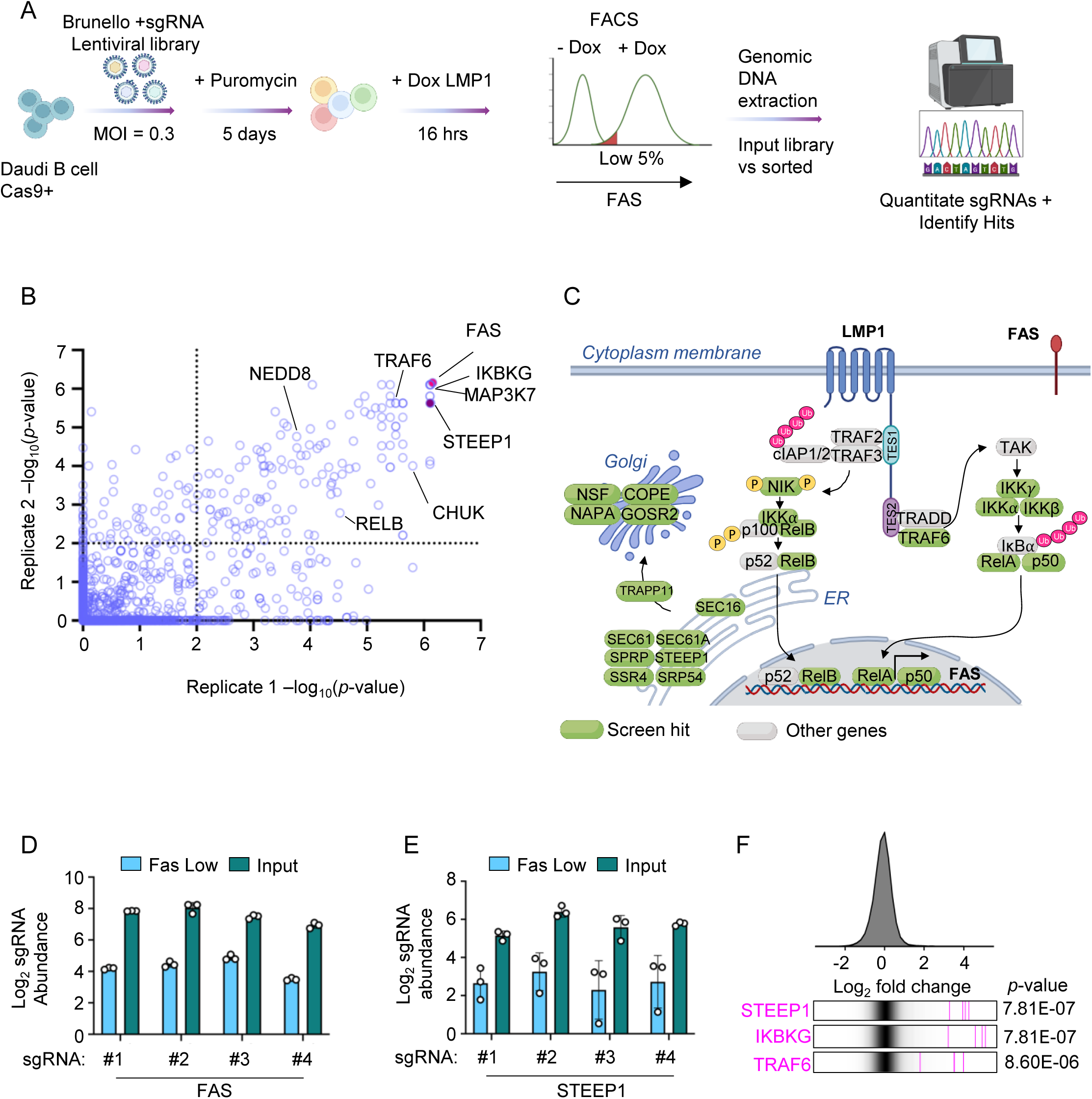
Genome-wide CRISPR/cas9 screen for LMP1 B cell dependency factors. (A) Human genome-wide CRISPR/Cas9 screen for LMP1 B cell dependency factors. Cas9+ Daudi B cells were transduced with the Brunello single guide RNA (sgRNA) library at a multiplicity of infection (MOI) of 0.3. Following transduced cell puromycin selection, LMP1 expression was induced by addition of doxycycline (200ng/mL) for 16 hours. The 5% of cells with the lowest expression of LMP1-induced target gene FAS were sorted. sgRNAs were PCR amplified from sorted vs input cells, and statistically significant hits were identified from n=3 biological screen replicates. (B) Scatter plot visualization of key hits from two screen replicates. STEEP1 was a top hit in both replicates shown. (C) Selected top hits highlighted in green. (D-E) Log2 sgRNA abundance of sgRNAs targeting *Fas* (D) and *STEEP1* (E) in the sorted 5% of cells with the lowest FAS expression (blue) versus unsorted input library (green). (F) Rug plots showing the distribution of Brunello human genome-wide CRISPR screen sgRNA log2 fold-change values from FAS^low^ sorted vs input cells. Values for sgRNAs targeting STEEP1, IKBKG (encodes NEMO) or TRAF6 (pink lines) are overlaid on gray gradients depicting all Brunello sgRNA library values. Average values from 3 screen biological replicates are shown.

Notably, we used FAS expression as a readout for our CD40 CRISPR screen [53, 54].

To screen for candidate LMP1 regulators, we used fluorescence-activated cell sorting (FACS) to sort for the 5% of cells with lowest plasma membrane (PM) FAS levels but that maintained a threshold amount of GFP, indicating that knockout of these hits did not simply block our conditional system or due to housekeeping effects on transcription or translation. The CRISPR screen was performed with biological triplicate replicates.

Genomic DNA was extracted from unsorted input and FACS-sorted cells, and PCR-amplified sgRNA abundance was quantified by next-generation DNA sequencing (**Figure 1A**). Hits, against whose sgRNA were enriched in sorted versus input populations, were identified by the STARS algorithm [60]. Applying a stringent false discovery rate threshold of q< 0.01, we identified 223 genes as putative positive regulators of LMP1 signaling, whose knockout diminished LMP1-mediated FAS induction, but not of our doxycycline-induced GFP expression control (**Figure 1B-C**, **Table 1**). Multiple well-established mediators of LMP1 signaling scored as hits, including genes encoding TRAF6, all three IKK complex subunits (IKKα, IKKβ and IKKγ), and the NF-κB transcription factor subunits p50 and p52 (**Figure 1B-C**).

Validating our experimental approach, *FAS* emerged as the most significantly enriched sgRNA target in cells sorted for low FAS expression, with sgRNAs targeting *FAS* ranked as the #3, 4, 5 and 21 most differentially expressed in sorted vs input populations out of the ∼76,000 sgRNAs in the Brunello library (**Figure 1B-F**). Intriguingly, multiple additional hits were not previously implicated in LMP1 biogenesis or signaling **(Table S1)**. Several of these have roles intracellular transport. These included three signal recognition particle (SRP) components and one subunit of the SRP receptor, likely reflecting their involvement in the endoplasmic reticulum membrane insertion of LMP1 and/or FAS. Five components of the COPII complex, which mediates ER-to-Golgi vesicular transport scored, as did the STING ER Exit Protein 1 (STEEP1), which regulates STING ER to Golgi trafficking [61] (**Figure 1C**, **Table S1**). sgRNAs targeting STEEP1 ranked at 55, 77, 81, and 144 most enriched in the sorted vs input population (**Fig. 1F**), suggestive of on-target CRISPR effects.

We next cross-compared results from the LMP1 screen with our published CD40 CRISPR screen [53] to define shared vs LMP1 selective host dependency factors (**Figure S1A**). Using an adjusted p-value< 0.05 cutoff, 35 genes were identified as important for both CD40 and LMP1-induced FAS expression (**Figure S1B**). These included genes encoding key regulators of NF-κB signaling: all three IKK complex subunits, the kinase TAK1, and the ubiquitin ligase FBXW11 that mediates ubiquitination of the canonical NF-κB pathway inhibitor IκBα (**Figure S1B, Table S1**). Notably, STEEP1 did not score in the CD40 screen (**Figure S1B**), suggesting that it is not important for either CD40 or FAS trafficking and underscoring that it does not chaperone all ER transmembrane protein cargo.

To validate STEEP1 as a hit, we treated CRISPR control or STEEP1 depleted cells with CD40 ligand (CD40L). As an orthogonal readout to FAS induction, we measured effects of STEEP1 depletion on processing of the p100 NF-κB transcription factor subunit to the active p52 form, which is stimulated by non-canonical NF-κB pathway activation.

CD40L treatment robustly induced p100:p52 processing in both control and STEEP1 depleted cells (**Figure S1C**), suggesting that STEEP1 is dispensable for CD40 trafficking to the plasma membrane and for CD40 signaling. Similarly, STEEP1 depletion did not significantly change control doxycycline-induced GFP expression or plasma membrane CD37 levels, further suggesting that STEEP1 does not generally regulate membrane protein trafficking out of the ER (**Figure S2**). By contrast, IKKγ or TRAF6 depletion strongly impaired LMP1-induced FAS expression (**Figure S2**).

STEEP1 depletion reduced both plasma membrane and intracellular FAS expression to a similar extent in LMP1+ Daudi and Akata B cells, further suggesting effects at the level of LMP1 signaling rather than FAS trafficking (**Figure 2**). Taken together, these results raise the possibility that STEEP1 supports trafficking of LMP1, but not all ER transmembrane protein cargo.

**Figure 2.**
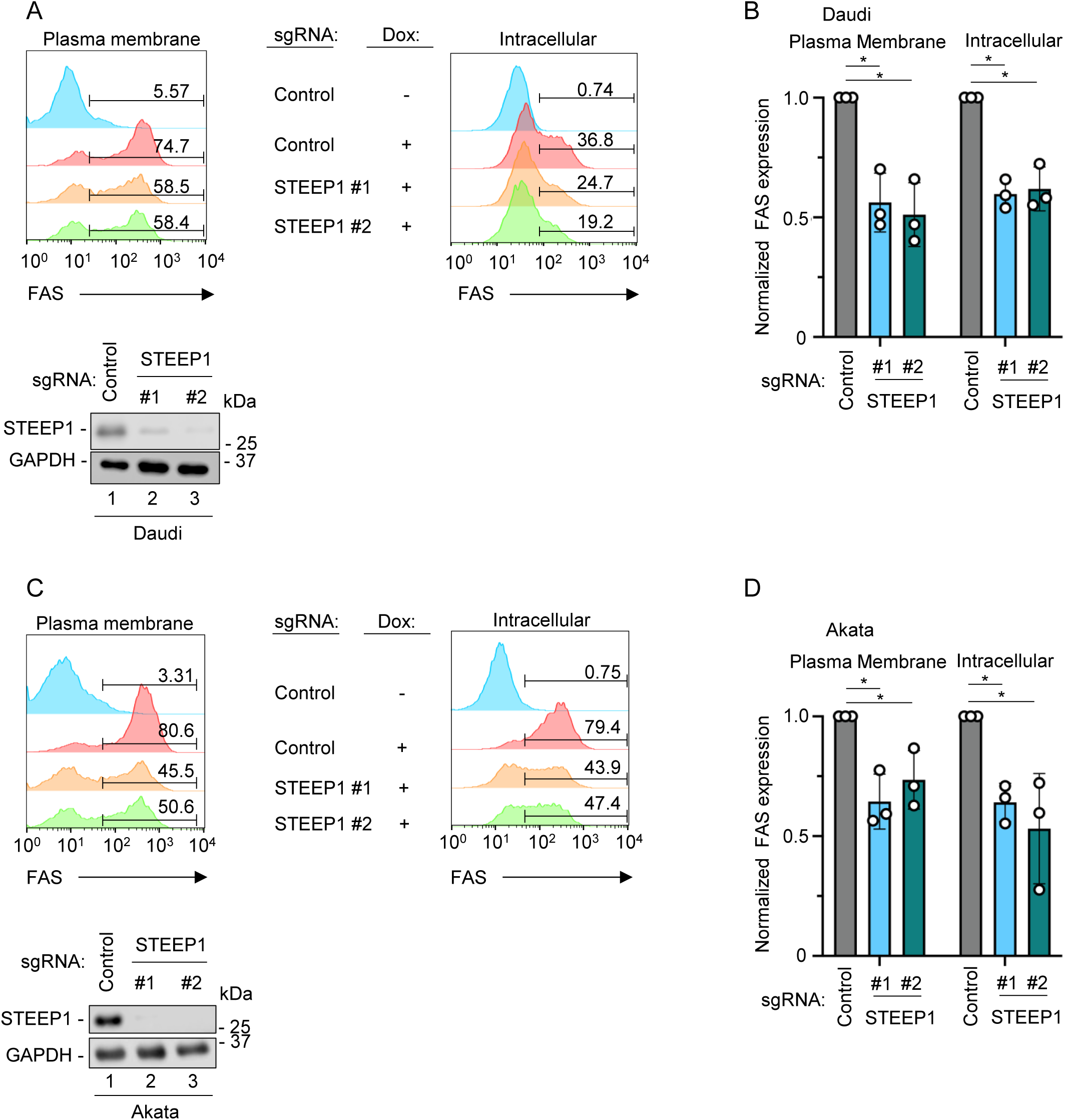
STEEP knockout impaired LMP1-mediated FAS upregulation. (A) FACS analysis of plasma membrane (left) and intracellular (right) FAS levels in Cas9+ Daudi cells with control or independent Brunello STEEP1 targeting sgRNAs and induced for LMP1 by 200ng/ml Dox for 16 hours, as indicated. Shown below, immunoblot of whole cell lysates (WCL) from Daudi cells expressing the indicated sgRNAs. (B) Normalized FAS plasma membrane vs intracellular mean fluorescence intensity (MFI) ± S.D. values from n=3 biological replicates as in (A). Values were normalized to control sgRNA levels. (C) FACS analysis of plasma membrane (left) and intracellular (right) FAS levels in Cas9+ Akata cells with control or independent Brunello STEEP1 targeting sgRNAs and induced for LMP1 by 200ng/ml Dox for 16 hours, as indicated. Shown below, immunoblot of whole cell lysates WCL from Akata cells expressing the indicated sgRNAs. (D) Normalized Akata FAS plasma membrane vs intracellular MFI ± S.D. values from n=3 biological replicates as in (C). Values were normalized to control sgRNA levels. Blots in A and C are representative of n=3 replicates.

### STEEP1 is a key host dependency factor for LMP1 signaling

We next characterized STEEP1 depletion effects on LMP1 signaling. We expressed control or independent Brunello library sgRNA targeting STEEP1 in either Cas9+ Daudi or Akata B cells and then induced LMP1 expression for 16 hours. STEEP1 depletion was validated by immunoblot (**Figure 3A-B**). In both Burkitt B cell contexts, STEEP1 depletion strongly impaired LMP1-driven p100:p52 processing indicative of non-canonical NF-κB activity, p38 phosphorylation indicative of MAP kinase activity, and LMP1 target gene FAS and TRAF1 induction (**Figure 3A-B**). To next examine STEEP1 depletion effects in the lymphoblastoid B-cell context, we expressed control or STEEP1 targeting sgRNA in GM12878 LCLs. Since LCLs are dependent on LMP1 signaling for survival, we measured depletion effects on LMP1 signaling at 5 days post-sgRNA expression. As observed in the Burkitt cell context, STEEP1 depletion impaired p100:p52 processing, p38 phosphorylation and LMP1-driven FAS and TRAF1 expression, indicative of an obligatory STEEP1 role in LMP1 signaling in LCLs (**Figure 3C-D**).

**Figure 3.**
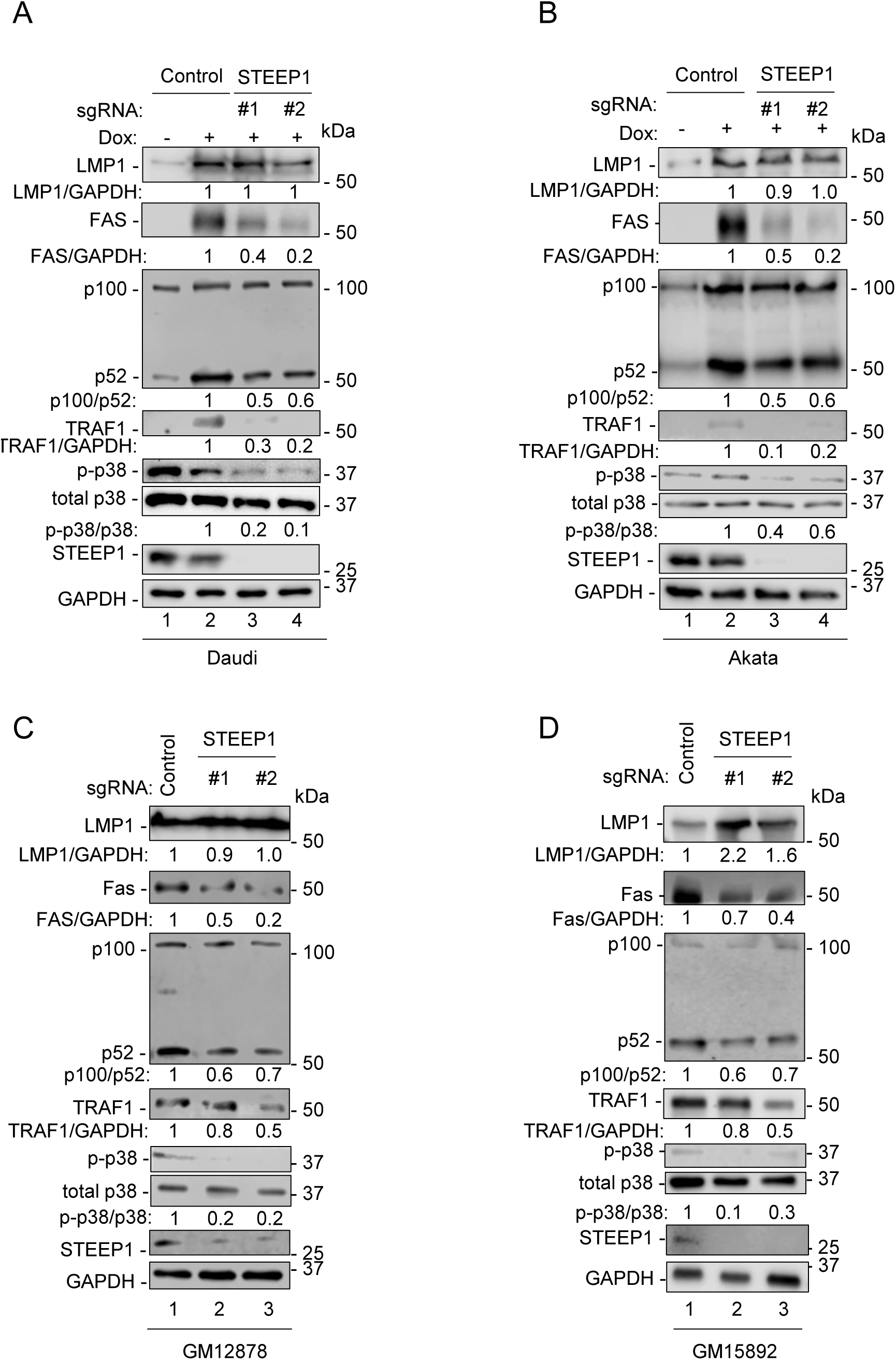
STEEP1 depletion impairs LMP1 signaling. (A) Analysis of STEEP1 depletion effects on LMP1 signaling in Daudi B cells. Immunoblot analysis of WCL from Cas9+ Daudi cells that expressed control or STEEP1 targeting sgRNAs and that were Dox (200ng/mL) induced for LMP1 expression for 16 hours. Densitometry indicates ratios as indicated and were normalized to sgRNA control levels. (B) Analysis of STEEP1 depletion effects on LMP1 signaling in Akata B cells. Immunoblot analysis of WCL from Cas9+ Akata cells that expressed control or STEEP1 targeting sgRNAs and that were Dox (200ng/mL) induced for LMP1 expression for 16 hours. (C) Analysis of STEEP1 depletion effects on LMP1 signaling in GM12878 LCLs. Immunoblot analysis of WCL from Cas9+ GM12878 that expressed control or STEEP1 targeting sgRNAs for 5 days. (D) Analysis of STEEP1 depletion effects on LMP1 signaling in GM15892 LCLs. Immunoblot analysis of WCL from Cas9+ GM15892 that expressed control or STEEP1 targeting sgRNAs for 5 days. Blots are representative of n=3 replicates.

Since LMP1 signaling is essential for LCL but not Burkitt B cell growth and survival, we next tested STEEP1 depletion effects on Burkitt versus LCL outgrowth. Control or STEEP1 targeting sgRNA were expressed in EBV-negative Daudi or Akata Burkitt cells, or in GM12878 or GM15892 LCLs. Live cell numbers were then measured for seven days. Interestingly, STEEP1 depletion rapidly and strongly blocked GM12878 and GM15892 proliferation. By contrast, it impaired Daudi but not Akata Burkitt B cell proliferation (**Figure 4**). These results are consistent with a model in which STEEP1 plays key roles as an LMP1 dependency factor in support of LCL growth and survival.

**Figure 4.**
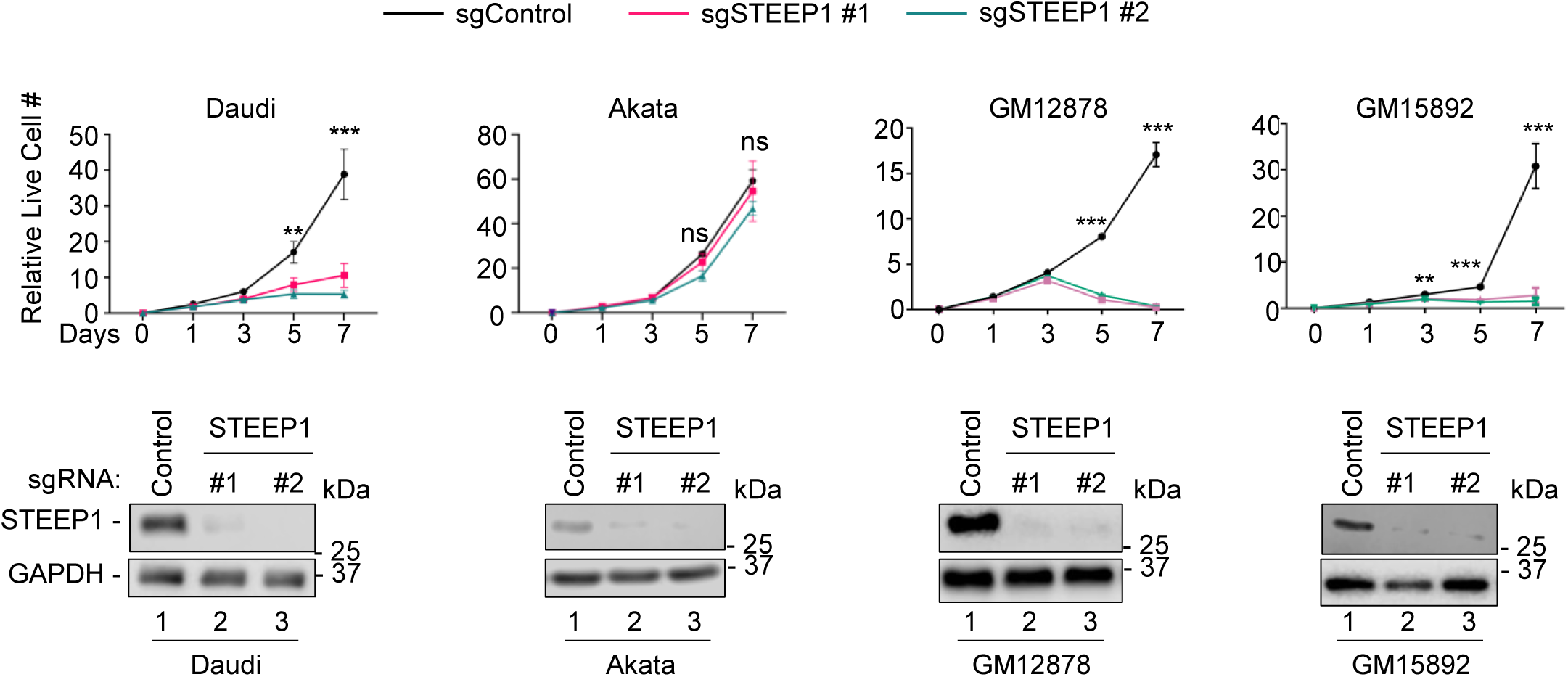
STEEP1 is a host dependency factor for LCL proliferation. STEEP1 effects on EBV transformed B cell proliferation. Shown are mean ± SD relative live cell values of the indicated Cas9+ B cell on the indicated days post-puromycin selection of cells transduced with lentivirus that express the indicated control or STEEP1 sgRNAs. Shown beneath each growth curve are immunoblots of WCL from each cell line, collected at day 3. **, p<0.01; ***, p<0.001. ns, non-significant.

### STEEP1 supports LMP1 trafficking

LMP1 signals from plasma membrane or intracellular signaling sites [16, 26, 31, 45–47]. To test the hypothesis that STEEP1 supports LMP1 trafficking, we analyzed LMP1 subcellular localization in control versus STEEP1 knockout cells using subcellular fractionation. For this analysis, we fractionated Daudi cell lysates by mechanical lysis to maintain the integrity of the membrane and avoid contaminating the membrane-bound species with the cytosolic compartments. CD9 was used as a plasma membrane marker, GM130 as a Golgi marker and GAPDH as a soluble protein marker. Daudi B cell STEEP1 depletion by either of two sgRNAs diminished the proportion of LMP1 that localized to the plasma membrane, as judged by immunoblot of fractionated samples (**Figure 5A-B**). Notably however, both sgRNAs depleted FAS expression, consistent with our CRISPR screen results. Likewise, STEEP1 depletion by either of two sgRNAs diminished LMP1 plasma membrane subcellular localization in GM12878 LCLs (**Figure 5C-D**).

**Figure 5.**
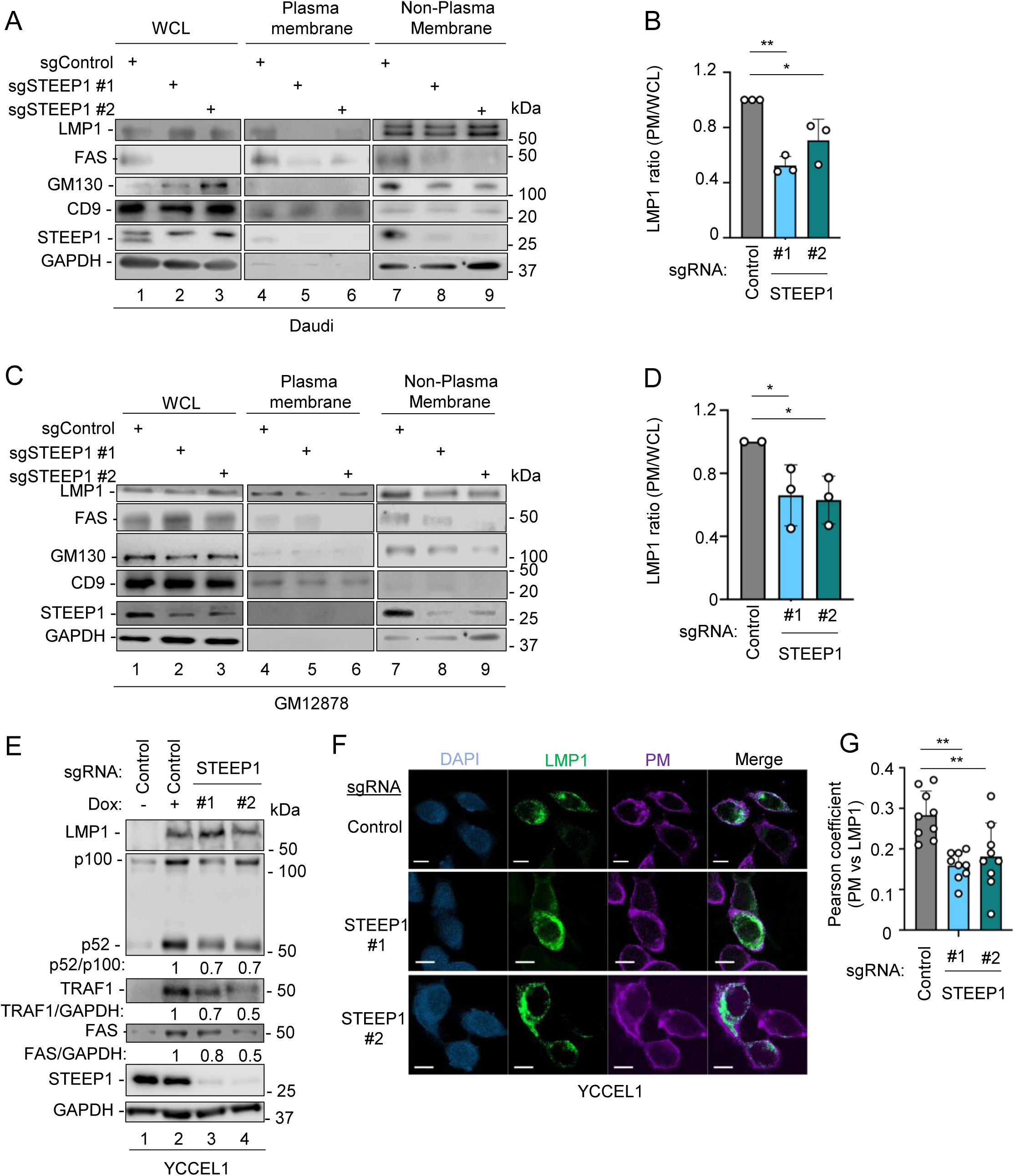
STEEP1 supports LMP1 plasma membrane trafficking in B and epithelial cells. (A) Effects of STEEP1 depletion on LMP1 plasma membrane subcellular localization in Daudi cells. Immunoblot analysis of extracts from Cas9+ Daudi cells that expressed control or STEEP1 sgRNAs and that were induced for LMP1 expression for 16 hours prior to preparation of WCL or of plasma membrane or non-plasma membrane fractions by centrifugation. GM130, CD9 and GAPDH were used as Golgi, plasma membrane and cytosolic markers, respectively. (B) Quantification of plasma membrane LMP1 to total cellular LMP1 ratio from 3 replicates as in (A). (C) Effects of STEEP1 depletion on LMP1 plasma membrane subcellular localization in GM21878 LCLs. Immunoblot analysis of extracts from Cas9+ GM12878 LCLs that were puromycin selected for three days following transduction with lentivirus expressing control or STEEP1 sgRNAs. (D) Quantification of plasma membrane LMP1 to total cellular LMP1 ratio from 3 replicates as in (C). (E) STEEP1 depletion effects on LMP1 signaling in YCCEL1 gastric carcinoma cells. Immunoblots of WCL from Cas9+ YCCEL1 expressing control or STEEP1 targeting sgRNA and induced for LMP1 by 200 ng/ml Dox for 16 hours. (F) STEEP1 depletion effects on LMP1 subcellular trafficking in YCCEL1. Confocal microscopy analyses of LMP1 (green) subcellular localization versus plasma membrane (PM, purple). Cells were treated as in (E) except for seeding onto coverslips 24 hours before Dox treatment. Scale bar=10 µm. (G) Quantification of STEEP1 depletion effects on LMP1 plasma membrane localization. Images as in (F) were analyzed by ImageJ using the colo2 plug-in. Pearson’s coefficient was calculated from n=10 LMP1+ cells. One-way ANOVA, *p<0.05 **, p<0.01. Blots in A, C and E are representative of n=3 independent replicates.

We next used human tumor derived EBV+ YCCEL1 gastric carcinoma cells to analyze STEEP1 depletion effects on LMP1 signaling and trafficking in an epithelial cell context. Since YCCEL1 have the EBV latency I program (EBNA1 only), we engineered doxycycline inducible LMP1 expression as well as stable Cas9 expression. STEEP1 depletion by either of two sgRNAs diminished LMP1-driven p100:p52 processing, TRAF1 and FAS target gene upregulation (**Figure 5E**). Whereas we observed substantial co-localization between LMP1 and an amine-reactive plasma membrane dye in cells with control sgRNA expression, STEEP1 depletion significantly altered LMP1 subcellular localization in YCCEL1 cells, as evidenced by diminished overlap between LMP1 and the plasma membrane signals (**Figure 5F-G**).

### STEEP1 promotes LMP1 trafficking from ER to Golgi

LMP1 signals from plasma membrane and intracellular membrane sites, but does not efficiently signal from the ER [47, 62]. Consistent with this, STEEP1 depletion by independent Brunello sgRNAs impaired LMP1 egress from the ER, as judged by increased overlap of LMP1 immunostain signals with that of the ER resident protein KDEL marker (**Figure 6A-B**). Also in support of an important STEEP1 role in LMP1 ER egress, STEEP1 depletion significantly diminished LMP1 trafficking to the Golgi, as judged by confiocal microscopy, where STEEP1 KO diminished overlap between LMP1 and Golgi marker GM130 signals (**Figure 6C-D**).

**Figure 6.**
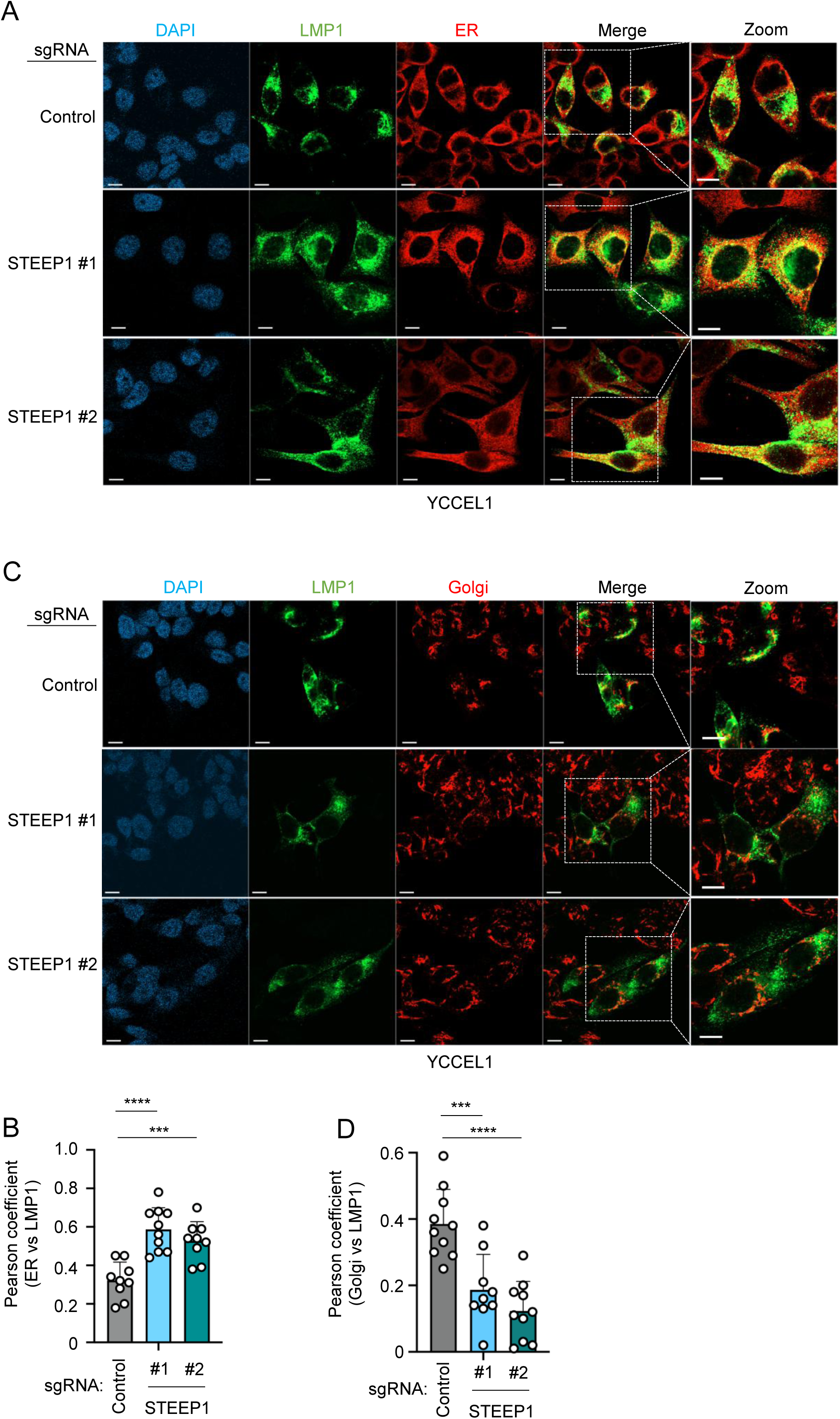
STEEP1 facilitates LMP1 trafficking from ER to Golgi. (A) Analysis of STEEP1 depletion effects on LMP1 ER egress. Confocal microscopy analysis of Cas9+ YCCEL1 cells with control or STEEP1 sgRNAs and Dox-induced for LMP1 expression for 16 hours. Shown are signals for LMP1-HA (green), the ER marker KDEL (red) and DAPI (blue). Scale bar=10 µm. (B) Quantification of LMP1/KDEL co-localization from cells as in (A). Images were analyzed by ImageJ using the colo2 plug-in. Pearson’s coefficient was calculated from n=10 cells. One-way ANOVA *** p<0,001, **** p<0.0001. (C) Analysis of STEEP1 depletion effects on LMP1 trafficking through the Golgi. Confocal microscopy analysis of Cas9+ YCCEL1 cells with control or STEEP1 sgRNAs and Dox-induced for LMP1 expression for 16 hours. Shown are signals for LMP1-HA (green), the ER marker, Golgi marker GM130 (red) and DAPI (blue). Scale bar=10 µm. (D) Quantification of LMP1/GM130 co-localization from cells as in (C). Images were analyzed by ImageJ using the colo2 plug-in. Pearson’s coefficient was calculated from n=10 cells. One-way ANOVA *** p<0,001, **** p<0.0001.

### LMP1 associates with STEEP1

STEEP1 associates with the DNA sensor STING and serves as a positive regulator of STING signaling by promoting its ER egress [61]. We therefore next tested the hypothesis that LMP1 and STEEP1 associate with one another. In support of this, conditionally expressed HA-tagged LMP1 co-immunoprecipitated STEEP1 from Daudi cell extracts (**Figure 7A**). To next test whether LMP1 associates with STEEP1 in LCLs, we utilized LCLs that express N-terminal FLAG-tagged LMP1 at physiological levels (the sequence encoding FLAG was knocked into the EBV genomic LMP1 locus [43]). Anti-FLAG-immunopurified LMP1, but not control IgG, co-immunoprecipitated STEEP1 (**Figure 7B**). We next tested whether STEEP1 could reciprocally co-immunoprecipitate LMP1 from LCL extracts. To do so, we stably expressed FLAG-tagged STEEP1 in GM12878 LCLs. anti-FLAG but not control IgG pulldown co-immunoprecipitated LMP1 (**Figure 7C**). In further agreement with the model that LMP1 and STEEP interact, we observed substantial overlap in confocal microscopy signals between LMP1 and STEEP1 and STEEP1 in YCCEL1, which were used rather than B-cells given their larger cell and cytoplasmic volumes to enhance microscopy visualization (**Figure 7D**).

**Figure 7.**
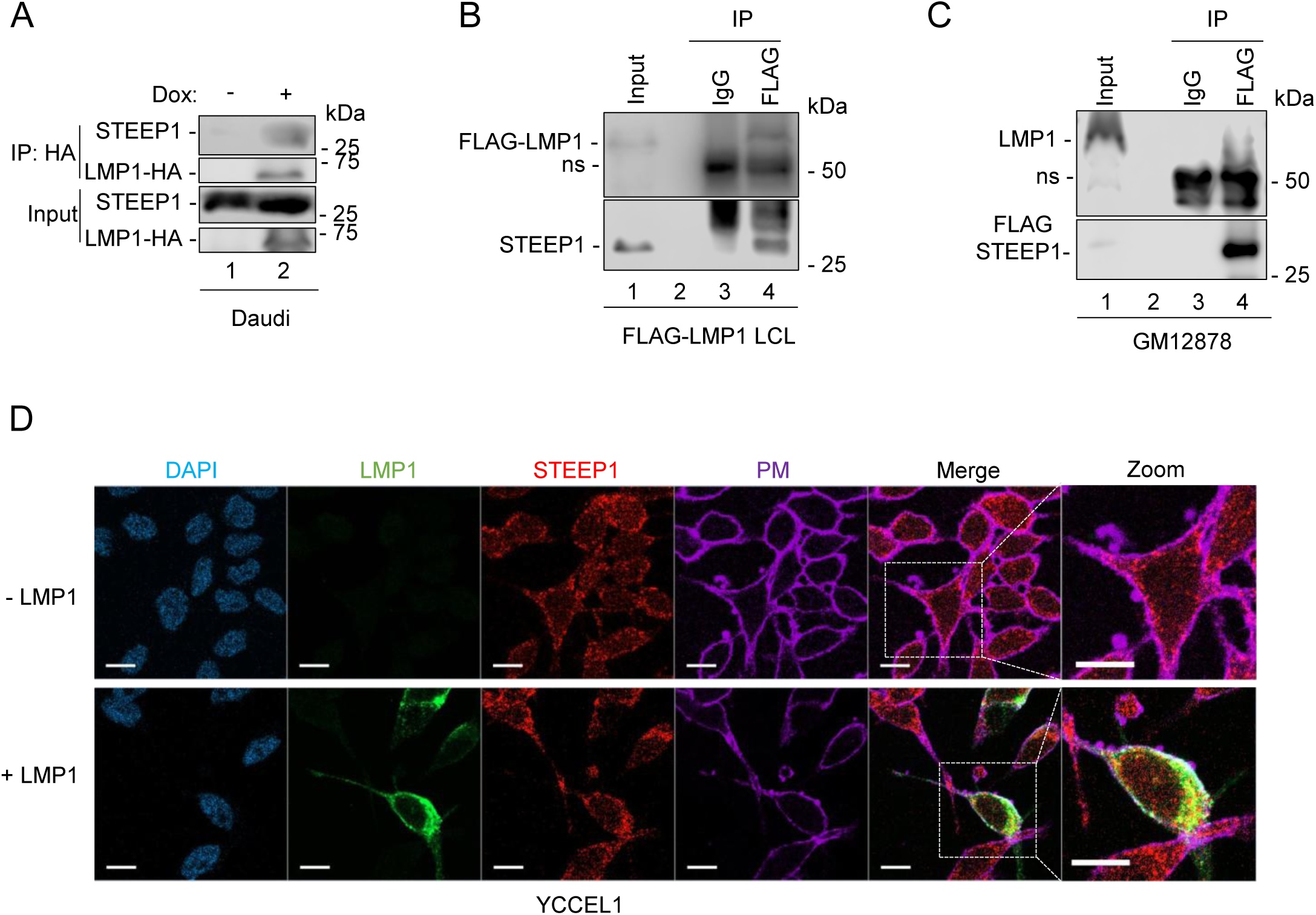
STEEP1 associates with LMP1. (A) Analysis of STEEP1/LMP1 association in Daudi B cells. 1% WCL vs anti-HA pulldown was subjected to immunoblot analysis. LMP1 was Dox-induced (200 ng/ml) for 16 hours prior to cell lysis. (B) Analysis of STEEP1/LMP1 association in FLAG-LMP1 LCLs. LCLs derived from EBV in which the FLAG epitope tag was knocked into the EBV genomic LMP1 locus were used. 1% WCL, control IgG or anti-FLAG immunoprecipitated complexes were subjected to immunoblot analysis, as indicated. ns, non-specific. (C) Analysis of STEEP1/LMP1 association in GM12878 LCLs. GM12878 1% WCL, control IgG or anti-FLAG immunoprecipitated complexes were subjected to immunoblot analysis, as indicated. ns, non-specific. (D) Confocal microscopy analysis of LMP1/STEEP1 association. Shown are images from YCCEL1 cells with stable FLAG-STEEP expression, mock induced or Dox-induced for LMP1 for 16 hours. Following PM staining (purple), cells were fixed/permeabilized and stained for FLAG (red), LMP1 (green) or DAPI (blue). Scale bar=10µm. Blots are representative of n=3 replicates.

### The LMP1 N-terminal cytoplasmic tail mediates association with STEEP1

Most known LMP1 interacting proteins associate with its C-terminal tail, though LMP1 transmembrane domains 3 and 4 are necessary for its association with PRA1, a host factor that supports LMP1 trafficking [62]. To identify LMP1 regions critical for its association with STEEP1, we expressed wildtype (WT) LMP1, or LMP1 deletion mutants lacking N-terminal cytoplasmic tail residues 6-17 or 18-24, or C-terminal tail deletion mutants lacking residues 231-286 or the entire tail (residues 187-386) (**Figure 8A**). Notably, residues 6-17 were previously identified as important for LMP1 subcellular trafficking to the plasma membrane and for EBV-mediated primary human B cell transformation [63].

**Figure 8.**
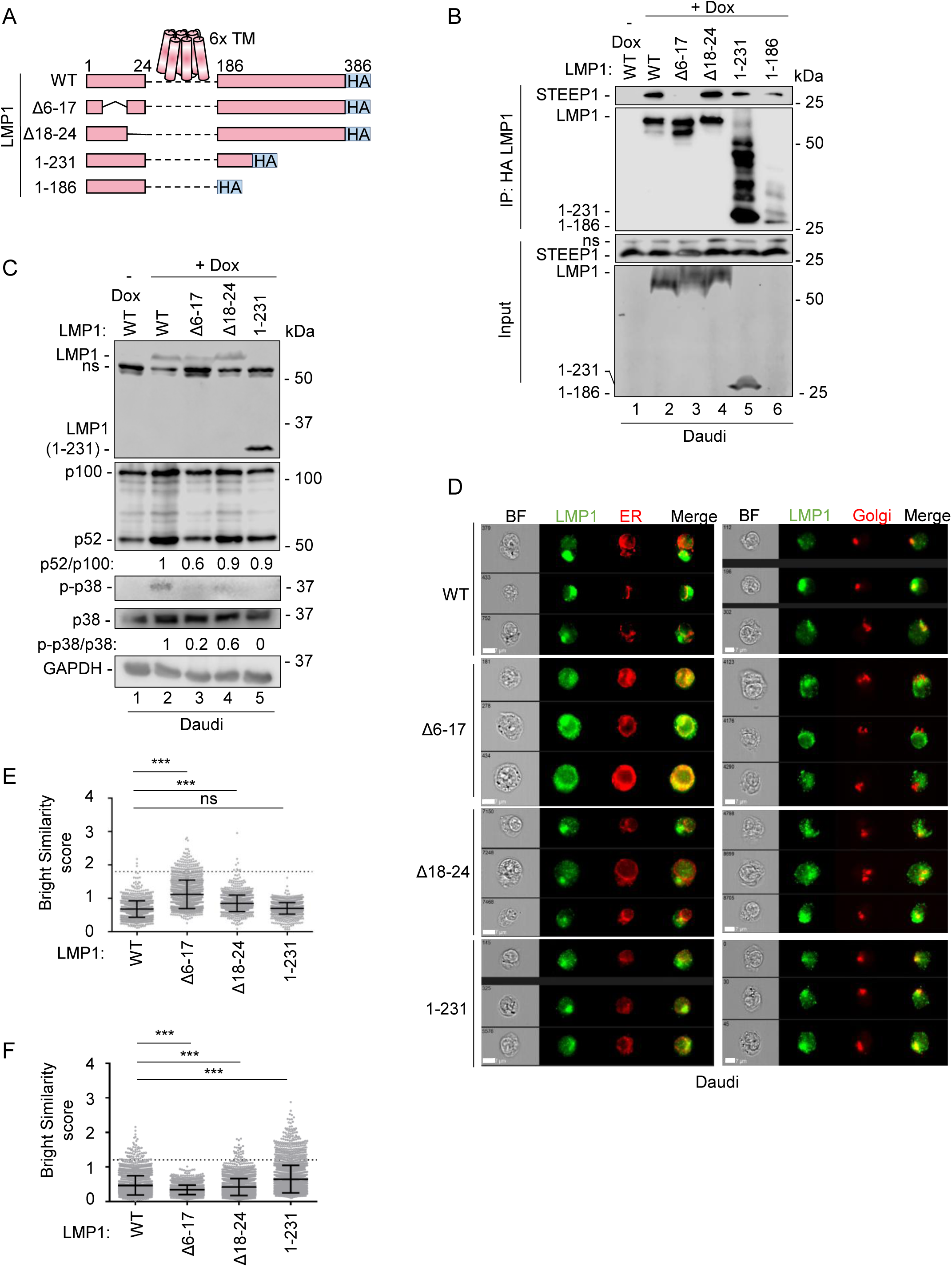
STEEP1 associates with the LMP1 N-terminal cytoplasmic tail. (A) Schematic model of C-terminal HA-tagged LMP1 constructs. TM, transmembrane domain. (B) Analysis of LMP1 N-terminal vs C-terminal tail regions important for STEEP1 association. Expression of the indicated LMP1 constructs were Dox-induced in Daudi cells for 16 hours. HA-immunoprecipitated complexes or 10% WCL were subject to immunoblot, as indicated. ns, non-specific. (C) Analysis of wildtype versus mutant LMP1 signaling. Immunoblot analysis of WCL from Daudi cells induced for the indicated LMP1 cDNA expression for 16 hours. (D) Analysis of LMP1 N or C-terminal tail roles in egress from the ER. Shown are ImageStream analyses of Daudi cells Dox-induced for expression of the indicted LMP1 cDNAs for 16 hours. Cells were stained for HA-LMP1 (green), KDEL (red) or GM130 (red). BF, brightfield. White bar=7µm. (E) Quantification of LMP1/KDEL co-localization. Live cells as in (D) were analyzed by IDEA software to yield a Bright Similarity Score R3. Bright Similarity Scores were analyzed in n=2500 LMP1+ cells. A value > 1.8 indicates colocalization. P-values were calculated by one-way ANOVA with Dunnett’s test and indicate comparison with WT LMP1+ cells. ***, p<0.0001, ns, non-significant. (F) Quantification of LMP1/GM130 co-localization. Live cells as in (D) were analyzed by IDEA software to yield a Bright Similarity Score R3. Bright Similarity Scores were analyzed in n=2500 LMP1+ cells. A value > 1.2 indicates colocalization. P-values were calculated by one-way ANOVA with Dunnett’s test and indicate comparison with WT LMP1+ cells. ***, p<0.0001, Dunnet’s test.

Deletion of LMP1 residues 6-17 strongly diminished LMP1 association with STEEP1, whereas deletion of residues 231-386 or the entire C-terminal tail did not significantly appreciably alter association with STEEP1 (**Figure 8B**). Signaling by LMP1 Δ6-17 was impaired, as judged by diminished p100:p52 processing and p38 phosphorylation relative to levels in Daudi cells with expression of wildtype LMP1, albeit with reduced steady state LMP1Δ6-17 levels (**Figure 8C**). Because the expression of the 1-186 mutant was lower compared to other mutants, we did not include it in our further tests. In Daudi cells, deletion of LMP1 residues 6-17 increased overlap between LMP1 and ER marker KDEL signals, but diminished overlap with the Golgi GM130 marker by ImageStream analysis (**Figures 8D-F**). Similarly, LMP1 Δ6-17 exhibited increased overlap with the ER KDEL marker in YCCEL1 (**Fig. S3**). Taken together, these results suggest that the LMP1 N-terminal tail mediates association with STEEP1 to support LMP1 ER egress (**Figure 9**).

**Figure 9.**
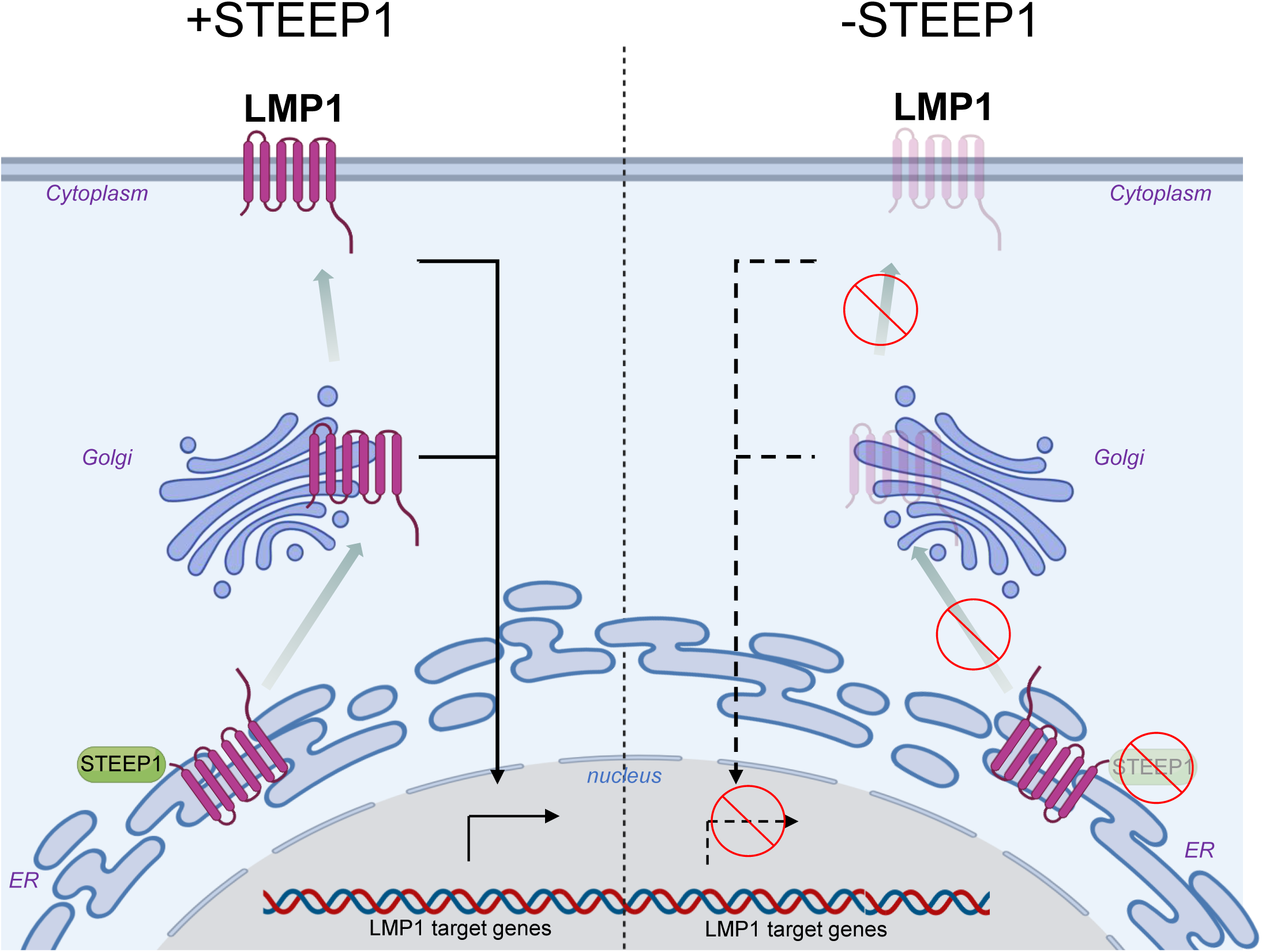
STEEP1 facilitates LMP1 trafficking from ER to Golgi. Schematic Mode. Left: STEEP1 associates with the N-terminal tail of ER-resident LMP1 molecules, and promotes their egress from the ER. Right: in the absence of STEEP1 or where LMP1 N-terminal tail/STEEP1 association is disrupted, LMP1 ER egress is perturbed, impairing its C-terminal tail signaling from distal membrane sites.

## Discussion

Here, we used a human genome-wide CRISPR-Cas9 screen to identify host factors important for LMP1 function. This revealed STEEP1 as a major host dependency factor that supports LMP1 signaling. STEEP1 depletion impaired signaling from both LMP1 TES1 and TES2 and rapidly diminished lymphoblastoid B-cell proliferation.

STEEP1/LMP1 association was dependent on the LMP1 N-terminal tail. STEEP1 depletion impaired LMP1 ER egress, as did deletion of the LMP1 N-terminal tail residues important for association with STEEP1. Together, these studies establish STEEP1 as an important host factor that supports LMP1 trafficking to cellular signaling sites.

Previous studies have highlighted that LMP1 traffics out of the ER to signal from membrane sites, including within the Golgi, plasma membrane and intracellular endosomes [28, 45, 47, 64–67]. Major LMP1 subpopulations co-localize with Golgi markers [62, 68] or reside within intracellular compartments, likely endosomes [47], though this can vary substantially between B and epithelial cells. Yet, host factors that regulate LMP1 egress from the ER have remained elusive. The host factor PRA1 was identified as an essential chaperone for LMP1 ER-to-Golgi trafficking [62] and also plays a role in Rab GTPase trafficking [69, 70]. Interestingly, PRA1/LMP1 interaction is dependent on LMP1 transmembranes 3 and 4. As our screen did not find PRA1 as a host dependency factor, it remains possible that PRA1 functions primarily in LMP1 epithelial cell trafficking, as prior PRA1 studies were not performed in B cells [62].

While not previously been implicated in LMP1 trafficking, STEEP1 was identified as a key factor that supports signaling by the innate immune DNA sensor STING [61]. Like LMP1, STING must traffic out of the ER to signal, and egress from the ER is the rate-limiting step in STING signaling [61]. Following STING activation by cGAMP and phosphorylation by TAK1 [61, 71], STEEP1 binds STING and recruits PI3K complex I components to the ER membrane [61]. This induces local phosphatidylinositol-3-phosphate (PtdIns(3)P) production for membrane remodeling and the establishment of ER exit sites, thereby enabling COPII-mediated transport of STING to the Golgi for downstream innate immune signaling [61]. Further studies are required to define whether LMP1 and STEEP1 together induce ER membrane curvature, as has been observed for STING following activation by cGAMP [61].

It will be of interest to determine whether LMP1 trafficking is also regulated by COPII-mediated export, and if so, whether host factors Sar1 and Sec24 that are implicated in STING transport together with STEEP1 support LMP1 egress. Relatedly, it will be of interest to determine whether induction of PtdINS(3)P is necessary for LMP1 ER egress, and if so, whether overexpression of the PtdIns(3)P hydrolase MTMR3 inhibits LMP1 trafficking and signaling. Importantly, STEEP1 is not required for all COPII-mediated trafficking from the ER, as for example vesicular stomatitis virus G protein COPII vesicle trafficking is not dependent on STEEP1. Likewise, we did not identify STEEP1 as a hit in our prior CRISPR screen for host dependency factors for CD40 [53], and STEEP1 depletion did not appreciably impair CD9 targeting to the plasma membrane.

While signaling by the LMP1 C-terminal tail has been extensively studied, comparatively little has remained known about LMP1 N-terminal tail roles. Importantly, molecular genetic analysis highlighted an important role for N-terminal tail residues 6-17 in LMP1 subcellular trafficking and in support of EBV-mediated B-cell growth transformation [63]. The authors concluded that the N-terminal tail was therefore probably important in anchoring the first LMP1 transmembrane domain in the plasma membrane, since deletion of residues 6-17 altered membrane localization. Rather, we now suggest that a key LMP1 N-terminal cytoplasmic role is to instead support LMP1 signaling to intracellular trafficking sites through association with STEEP1 and activation of STEEP1 ER transport roles. As ER signals from post-ER membrane sites, we suggest that a key LMP1 N-terminal tail role is to subvert STEEP1 in support of LMP1 trafficking.

LMP1 is required for EBV-mediated B-cell transformation, for the growth and survival of lymphoblastoid B-cells, and is also implicated in epithelial cell transformation with roles in nasopharyngeal carcinoma pathogenesis [8, 19, 23, 34, 37, 40, 52, 72–75].

Recently, LMP1 signaling has also been implicated in multiple sclerosis pathogenesis [14, 15]. These observations highlight LMP1 as an attractive therapeutic target, though LMP1 itself has not yet proven to be druggable. Our study suggests that targeting STEEP1, or STEEP1/LMP1 interaction, may instead be an attractive target to block LMP1 ER egress and signaling. Interestingly, CRISPR dependency map (DepMap.org) indicates that STEEP1 knockout is tolerated by most of the nearly 700 cell lines tested. The exquisite LCL dependence on STEEP1 is therefore notable and further highlights STEEP1 as a potential therapeutic target. It will therefore be of interest to develop and test inhibitors of STEEP1, or of the STEEP1/LMP1 N-terminal tail association, on EBV-driven B-cell immortalization and on proliferation of B and epithelial cell malignancies dependent on LMP1.

Several key questions regarding STEEP1 dependency factor roles in support of LMP1 remain to be addressed. First, it will be of interest to define STEEP1 residues necessary for interaction with LMP1, and whether these differ from residues that interact with STING. Second, it will be of interest to define whether STEEP1 depletion blocks EBV-mediated primary B-cell immortalization, and if so, whether this correlates with time-points at which LMP1 signaling becomes necessary for infected cell growth and survival. Third, it will be of interest to define whether the LMP1/STEEP1 interaction supports growth and survival of nasopharyngeal carcinoma or T and NK cell lymphoma cells, both of which can express LMP1. Fourth, it will be of interest to define whether LMP1 competes with STING for association with STEEP1 in a manner that impairs STING egress and signaling for innate immune evasion.

In conclusion, CRISPR-Cas9 genetic analysis highlighted STEEP1 as a host dependency factor important for LMP1 signaling. STEEP1 associates with the LMP1 N-terminal tail and supports LMP1 ER egress to intracellular signaling sites.

STEEP1/LMP1 interaction may therefore serve as a highly selective therapeutic target to block LMP1 signaling in the setting of malignancy or autoimmunity.

## Materials and methods

### Cell Culture

All cell lines, including EBV-negative Akata and Daudi Burkitt B cells, the LCLs GM15892 and GM12878, and the EBV+ gastric carcinoma YCCEL1 were maintained in a humidified incubator at 37∘C with 5% CO_2_. Cells were cultured in RPMI 1640 medium (Gibco) supplemented with 10% Fetal Bovine Serum (FBS) (Sigma-Aldrich), 2 mM L-glutamine, and 100 U/mL penicillin/streptomycin. Dox-inducible LMP1 cells lines were generated by lentiviral transduction using the pLIX-402 vector, as previously described [19, 76]. Lentivirus with Dox-inducible GFP expression were generated using the p-Inducer vector (Addgene plasmid #44012, a kind gift from Steven Elledge, [77]) and used to transduce the Dox-inducible LMP1 cells, which were selected by G418. In this manner, Dox treatment caused cells to express both LMP1 and a GFP control. For induction, cells were treated with 200 ng/mL dox.

### Genome-wide CRISPR/Cas9 screen

A human genome-wide CRISPR-Cas9 screen was conducted as previously described [53]. 130 million Cas9+ Daudi Dox-inducible LMP1 B cells were transduced with the Brunello sgRNA library at a multiplicity of infection (MOI) of 0.3 via spinoculation at 300x g for 2 hours in the presence of 4 μg/mL polybrene. Cells were placed in the incubator for 6 hours at 37°C and 5% CO_2_, pelleted and media was exchanged for fresh RPMI/10% FCS in the absence of polybrene. Libraries were prepared in biological triplicate. Successfully transduced cells were selected with 3 μg/mL puromycin 2 days post transduction for 48 hours. Each library was passaged every 72 hours, maintaining a minimum cell number of 40 million per replicate to ensure adequate library complexity. Five days after puromycin selection, a baseline (input) genomic DNA (gDNA) sample was extracted from 40 million cells per each of the three replicates, using the QIAGEN Blood and Cell Culture DNA Maxi Kit. Separately, 160 million cells from the library were treated with 200 ng/mL Dox for 16 hours to induce LMP1 expression. pINDUCER20-GFP plasmid where cells will express GFP once receiving Doxycycline. Cells were stained with APC-conjugated anti-FAS antibody (Biolegend #305612) and subjected to fluorescence-activated cell sorting (FACS) at the Brigham and Women’s Hospital Human Immunology Center. The 5% of cells with the lowest FAS expression but that expressed the Dox-induced GFP control were collected. Genomic DNA was extracted from sorted cells from each of three biological screen replicates, using the DNeasy Blood & Cell Culture DNA Kit (Qiagen # 13362). PCR-amplifed sgRNA abundances from input versus sorted cells was quantified by next-generation DNA sequencing [78] (Novogene). Screen hits were identified by the STARs algorithm [60], using an adjusted P-value cutoff of <0.05.

### CRISPR editing

CRISPR/Cas9-mediated gene editing was carried out in B-cell lines with stable Cas9 expression following established protocols [53, 60]. Control or gene-targeting sgRNAs from the Broad Institute Brunello libraries (see **Table S3** for sgRNAs and plasmids) were cloned into the lentiviral sgRNA expression vector with puromycin selection marker.

Correct sgRNA insertion was verified by Sanger sequencing. Lentiviral particles were produced by co-transfecting 293T cells with the sgRNA vector alongside VSV-G envelope and packaging plasmids using TransIT-LT1 reagent (Mirus Bio). Viral supernatants collected at 48 and 72 hours post-transfection were applied directly to target B cells. A GFP-targeting sgRNA served as a transduction control. Successfully transduced cells were selected using puromycin (3 μg/mL), and knockout efficiency was confirmed by immunoblotting.

### Immunoblot Analysis

Live cell numbers were normalized by the CellTiter-Glo (CTG) luminescence assay (Promega). Cells were lysed in RIPA buffer (Thermo Fisher). Equal amounts of cell lysates were separated by SDS-PAGE and transferred to nitrocellulose membranes. Membranes were blocked with 5% non-fat dry milk in TBST and probed overnight at 4°C with primary antibodies in TBST. The following primary antibodies were used: anti-LMP1 (OT22CN, a gift from Jaap Middeldorp), anti-STEEP1 (CST, #35136), anti-phospho-p38 (Thr180/Tyr182) (CST, # 9211S), Anti-total p38 (CST, # 8690s), anti-TRAF1 (CST, # 4710T), anti-p100/52 (Millipore Sigma # 05-361), anti-FAS (CD95) (CST, # 8023S), anti-GAPDH (EMD Millipore, # MAB374). HRP-conjugated secondary antibodies (CST, # 7074V) were used for detection, and signals were visualized using enhanced chemiluminescence (ECL) substrate (Bio-Rad). Densitometric analysis was performed using a LI-COR Odessey.

### Subcellular fractionation

10 million cells were subjected to subcellular fractionation, as previously described [61, 79]. Briefly, cells were mechanically lysed using a 27G needle. Loss of cell integrity was confirmed by microscopic analysis. Lysates were centrifuged at 700 xg for 10 min at 4°C to remove intact cells and nuclei. The supernatant was decanted and centrifuged for another 45 min at 21,000 x g. The pellet was washed once with lysis buffer. Equivalent protein amounts from the whole cell lysate (WCL) and the membrane fraction were analyzed via immunoblot. Fraction purity was validated using the following markers: CD9 (plasma membrane), GM130 (Golgi membrane) and GAPDH (cytosol). The fraction of plasma membrane localized LMP1 was quantitated by calculating LMP1 band intensity in the plasma membrane fraction to LMP1 intensity in the whole cell lysate fraction.

### Immunofluorescence and Confocal Microscopy

Plasma membranes were stained by the dye BriteMem following the manufacturer’s instructions (Biotium, #30098-T). Stained cells were then fixed with 4% paraformaldehyde for 10 minutes at room temperature, permeabilized with 0.5% Triton X-100 for 5 min at room temperature and blocked with 1% BSA in PBS for one hour at room temperature. Cells were then stained overnight at 4°C with primary antibodies targeting LMP1 (Abcam. #ab78113), mouse anti-HA and/or the following organelle markers: the ER marker KDEL (ThermoFisher cat# PA1-013) or the Golgi marker GM130 (CST, cat# 12480T) (1:400 dilution). Cells were then stained with Alexa Fluor 488 or 568 conjugated secondary antibodies (Invitrogen, # A-11008 or A-11031) in PBS (1: 1,000 dilution) for 1 hour at 37°C. Nuclei were counterstained with DAPI (ThermoFisher, # P36930). Images were acquired using a Zeiss confocal microscope with a 63x oil-immersion objective. Image analysis was performed as follows: colocalization analysis was performed using the Fiji (ImageJ) software platform.

Pearson’s Correlation Coefficients were calculated from at least 10 cells per condition across three independent experiments to quantify the degree of overlap between the LMP1 signal and KDEL or Golgi organelle markers.

### Immunoprecipitation

Immunoprecipitation was performed as previous described [80]. Briefly, cells were lysed in lysis buffer (50 mM Tris-HCl, pH 7.4, 150 mM NaCl, 1 mM EDTA, 1% NP-40) supplemented with protease inhibitors (Sigma, # 11836170001). If not stated specifically, usually 10% (v/v) of an aliquot from each lysate was preserved as the input. The rest of lysates were incubated with 2 μg of either control IgG, Anti-FLAG (Sigma #F3165), or Anti-HA magnetic beads (20 μL, ThermoFisher # PI88837) overnight at 4°C. For Anti-FLAG antibodies, antibody-protein complexes were captured using Protein A/G magnetic beads by incubating 2 hours (Pierce, #88803). Beads were washed extensively with PBST three times, and bound proteins were eluted by boiling in SDS loading buffer for 5 minutes.

### ImageStream

Imaging flow cytometry was performed on an ImageStream X [Mark II] (Luminex/Amnis) at 40x magnification. A minimum of 5,000-10,000 single cells were acquired per sample. Single-color compensation controls were prepared for each fluorochrome. Data analysis was performed using IDEAS software version [6.2]. Focused cells were gated using the Gradient RMS feature to exclude out-of-focus events. Debris and doublets were excluded based on Area and Aspect Ratio morphological features. Dead cells were gated out based on positive signals shown on the Zombie Aqua dye assigned to channel 11 (642 – 740 nm). LMP1-expressing cells were identified based on Alexa Fluorophore 488 positivity.

Colocalization between LMP1 and each subcellular compartment marker (KDEL for ER, GM130 for Golgi, and the PM stain) was quantified using the Bright Detail Similarity (BDS) R3 score. The BDS R3 score calculates the log-transformed Pearson correlation coefficient of bright punctate features within a 3-pixel radius mask applied to paired fluorescent channels on a cell-by-cell basis, with higher scores indicating greater spatial colocalization. A BDS R3 threshold of 1.8 for ER and 1.2 for Golgi was determined based on single-stained controls to define colocalized versus non-colocalized populations. The percentage of LMP1-expressing cells with BDS R3 scores above threshold was compared across wildtype LMP1 and each truncation mutant to determine the effect of N-terminal deletions on LMP1 subcellular localization.

### Cell Proliferation Assay

Cell growth was monitored by live cell number quantification using the Cell Titer Glo assay. Cells were plated in triplicate in 24-well plates after 2 days of puromycin selection at a density of 3×10^5^ cells/mL. Luminescence readings were taken at Days 0, 3, 5 and 7 days following puromycin plating. Results were normalized to the t=0 reading to determine the relative number of cells.

## Statistical Analysis

All quantitative data are presented as the mean ± Standard deviation of the Mean (SD) from at least three independent biological replicates. Statistical significance was determined using a two-tailed Student’s t-test or One-way ANOVA with appropriate post-hoc analysis (e.g., Dunnett’s multiple comparisons test), where applicable. A p-value of less than 0.05 (p<0.05) was considered statistically significant. Statistical analyses were performed using GraphPad Prism software (version 9). For statistical analyses for ImageStream, BDS R3 scores between wildtype LMP1 and truncation mutants were assessed using one-way ANOVA with Kruskal-Wallis test with Dunn’s correction.

## Acknowledgements

This work was supported by NIH R01 CA228700, U01CA275301 and P01 CA269043 to BEG. SL was supported by NIH P30 AI060354 Harvard University Center for AIDS Research (CFAR), which is supported by the following NIH Co-Funding and Participating Institutes and Centers: NIAID, NCI, NICHD, NHLBI, NIDA, NIMH, NIA, NIDDK, NIMHD, NIDCR, NINR, OAR, and FIC. YS, EMB, and BM were supported by American Cancer Society Post-doctoral Fellowships PF-23-1144614-01-IBCD, PF-23-898493-01-TBE, and PF-24-1250090-01-IBCD, respectively. The authors also acknowledge generous philanthropic support of George and Sandra K. Schussel. The funders had no role in study design, data collection and interpretation, or the decision to submit the work for publication. We thank Jaap Middeldorp for the generous gift of anti-LMP1 monoclonal antibody OT22CN.

## Supplementary Figure Legends

**Figure S1. STEEP1 is a host cell dependency factor for LMP1 signaling.**

(A) Schematic diagram cross-comparing LMP1 vs B cell co-receptor CD40 signaling. LMP1 and CD40 each activate NF-kB, JNK and JAK/STAT pathways amongst others.

(B) Cross-comparison of human genome-wide CRISPR/Cas9 screens for B cell factors that support LMP1 vs CD40 signaling. Scatter plot shows –log10(p-value) of hits from the LMP1 screen (see Fig. 1) vs our prior screen of CD40 signaling [53]. Both screens used Daudi cells and used FACsorting of cells with the lowest 5% of plasma membrane FAS as a readout. STEEP1 scored highly in the LMP1 but not CD40 screen, indicating that it is not required for CD40 signaling or for CD40 or FAS trafficking to the plasma membrane.

(C) STEEP1 depletion does not impair CD40 signaling. Immunoblot analysis of WCL from Cas9+ Daudi B cells expressing control or STEEP1 sgRNAs and stimulated by CD40L (50 ng/mL) for 48 hours as indicated. Blots represent n=3 independent biological replicates. P100:p52 ratios, indicative of non-canonical NF-kB activity, are shown.

**Figure S2. FACS analysis of control and screen hit sgRNAs.**

(A) STEEP1 depletion does not alter CD37 plasma membrane trafficking. FACS analysis of plasma membrane CD37 expression Cas9+ Daudi cells expressing control or STEEP1 sgRNAs and Dox-induced for LMP1 expression for 16 hours as indicated.

(B) STEEP1 depletion does not alter Dox-induced GFP reporter expression. FACS analysis of GFP expression in Cas9+ Daudi cells expressing control or STEEP1 sgRNAs and Dox-induced for control GFP expression for 16 hours. The same system that was used elsewhere for Dox-induced LMP1 expression was used here to drive GFP expression as a control.

(C) NEMO depletion strongly impairs LMP1-driven FAS expression. FACS analysis of plasma membrane FAS expression in Cas9+ Daudi cells expressing control or IKBKG (encodes NEMO) targeting sgRNAs and Dox-induced for LMP1 expression for 16 hours.

(D) TRAF6 depletion strongly impairs LMP1-driven FAS expression. FACS analysis of plasma membrane FAS expression in Cas9+ Daudi cells expressing control or TRAF6 targeting sgRNAs and Dox-induced for LMP1 expression for 16 hours.

**Figure S3. LMP1 egress from the ER is supported by its N-terminal cytoplasmic tail.**

(A) Analysis of LMP1 N vs C terminal cytoplasmic tail deletions on ER egress. Confocal microscopy analysis of YCCEL1 cells, Dox-induced for expression of the indicated LMP1 cDNA for 16 hours. Shown are signals for LMP1-HA (green), the ER marker KDEL (red), plasma membrane (purple) and DAPI (blue). Scale bar=10 µm.

(B) Colocalization analyses of ER and LMP1 (WT or N-terminal truncation mutants) in YCCEL1 cells as in (A). Images from n=5 cells were analyzed by ImageJ using the colo2 plug-in. Pearson coefficient was calculated from each cell. One-way ANOVA, *p<0.05, ns, non-significant.

## References

1. Damania, B., S.C. Kenney, and N. Raab-Traub, Epstein-Barr virus: Biology and clinical disease. Cell, 2022. 185(20): p. 3652–3670.

2. Frappier, L., Epstein-Barr virus: Current questions and challenges. Tumour Virus Res, 2021. 12: p. 200218.

3. Farrell, P.J., Epstein-Barr Virus and Cancer. Annu Rev Pathol, 2019. 14: p. 29–53.

4. Chiu, Y.F., K. Ponlachantra, and B. Sugden, How Epstein Barr Virus Causes Lymphomas. Viruses, 2024. 16(11).

5. Dawson, C.W., R.J. Port, and L.S. Young, The role of the EBV-encoded latent membrane proteins LMP1 and LMP2 in the pathogenesis of nasopharyngeal carcinoma (NPC). Semin Cancer Biol, 2012. 22(2): p. 144–53.

6. Healy, J.A. and S.S. Dave, The Role of EBV in the Pathogenesis of Diffuse Large B Cell Lymphoma. Curr Top Microbiol Immunol, 2015. 390(Pt 1): p. 315–37.

7. Shannon-Lowe, C., A.B. Rickinson, and A.I. Bell, Epstein-Barr virus-associated lymphomas. Philos Trans R Soc Lond B Biol Sci, 2017. 372 (1732).

8. Kieser, A., The Latent Membrane Protein 1 (LMP1): Biological Functions and Molecular Mechanisms. Curr Top Microbiol Immunol, 2025.

9. Wang, L.W., S. Jiang, and B.E. Gewurz, Epstein-Barr Virus LMP1-Mediated Oncogenicity. J Virol, 2017. 91(21).

10. Robinson, W.H., et al., Epstein-Barr virus as a potentiator of autoimmune diseases. Nat Rev Rheumatol, 2024. 20(11): p. 729–740.

11. Younis, S., et al., Epstein-Barr virus reprograms autoreactive B cells as antigen-presenting cells in systemic lupus erythematosus. Sci Transl Med, 2025. 17(824): p. eady0210.

12. Harley, J.B., et al., Transcription factors operate across disease loci, with EBNA2 implicated in autoimmunity. Nat Genet, 2018. 50(5): p. 699–707.

13. James, J.A., et al., An increased prevalence of Epstein-Barr virus infection in young patients suggests a possible etiology for systemic lupus erythematosus. J Clin Invest, 1997. 100(12): p. 3019–26.

14. Orr, N. and L. Steinman, Epstein-Barr virus and the immune microenvironment in multiple sclerosis: Insights from high-dimensional brain tissue imaging. Proc Natl Acad Sci U S A, 2025. 122(11): p. e2425670122.

15. Steinman, L. and S.S. Zamvil, Discoveries beyond molecular mimicry describe how EBV drives multiple sclerosis. Cell, 2026. 189(2): p. 345–347.

16. Kieser, A. and K.R. Sterz, The Latent Membrane Protein 1 (LMP1). Curr Top Microbiol Immunol, 2015. 391: p. 119–49.

17. Stunz, L.L. and G.A. Bishop, Latent membrane protein 1 and the B lymphocyte-a complex relationship. Crit Rev Immunol, 2014. 34(3): p. 177–98.

18. Kaye, K.M., et al., The Epstein-Barr virus LMP1 cytoplasmic carboxy terminus is essential for B-lymphocyte transformation; fibroblast cocultivation complements a critical function within the terminal 155 residues. J Virol, 1995. 69(2): p. 675–83.

19. Mitra, B., et al., Characterization of target gene regulation by the two Epstein-Barr virus oncogene LMP1 domains essential for B-cell transformation. mBio, 2023. 14(6): p. e0233823.

20. Price, A.M. and M.A. Luftig, To be or not IIb: a multi-step process for Epstein-Barr virus latency establishment and consequences for B cell tumorigenesis. PLoS Pathog, 2015. 11(3): p. e1004656.

21. Baichwal, V.R. and B. Sugden, Transformation of Balb 3T3 cells by the BNLF-1 gene of Epstein-Barr virus. Oncogene, 1988. 2(5): p. 461–7.

22. Moorthy, R.K. and D.A. Thorley-Lawson, All three domains of the Epstein-Barr virus-encoded latent membrane protein LMP-1 are required for transformation of rat-1 fibroblasts. J Virol, 1993. 67(3): p. 1638–46.

23. Wang, D., D. Liebowitz, and E. Kieff, An EBV membrane protein expressed in immortalized lymphocytes transforms established rodent cells. Cell, 1985. 43(3 Pt 2): p. 831–40.

24. Gires, O., et al., Latent membrane protein 1 of Epstein-Barr virus mimics a constitutively active receptor molecule. EMBO J, 1997. 16(20): p. 6131–40.

25. Eliopoulos, A.G., et al., The oncogenic protein kinase Tpl-2/Cot contributes to Epstein-Barr virus-encoded latent infection membrane protein 1-induced NF-kappaB signaling downstream of TRAF2. J Virol, 2002. 76(9): p. 4567–79.

26. Higuchi, M., K.M. Izumi, and E. Kieff, Epstein-Barr virus latent-infection membrane proteins are palmitoylated and raft-associated: protein 1 binds to the cytoskeleton through TNF receptor cytoplasmic factors. Proc Natl Acad Sci U S A, 2001. 98(8): p. 4675–80.

27. Coffin, W.F., 3rd, T.R. Geiger, and J.M. Martin, Transmembrane domains 1 and 2 of the latent membrane protein 1 of Epstein-Barr virus contain a lipid raft targeting signal and play a critical role in cytostasis. J Virol, 2003. 77(6): p. 3749–58.

28. Yasui, T., et al., Latent infection membrane protein transmembrane FWLY is critical for intermolecular interaction, raft localization, and signaling. Proc Natl Acad Sci U S A, 2004. 101(1): p. 278–83.

29. Kaykas, A., K. Worringer, and B. Sugden, LMP-1’s transmembrane domains encode multiple functions required for LMP-1’s efficient signaling. J Virol, 2002. 76(22): p. 11551–60.

30. Lee, J. and B. Sugden, A membrane leucine heptad contributes to trafficking, signaling, and transformation by latent membrane protein 1. J Virol, 2007. 81(17): p. 9121–30.

31. Huang, J., et al., Assembly and activation of EBV latent membrane protein 1. Cell, 2024. 187(18): p. 4996–5009 e14.

32. Fish, K., et al., Rewiring of B cell receptor signaling by Epstein-Barr virus LMP2A. Proc Natl Acad Sci U S A, 2020. 117(42): p. 26318–26327.

33. Uchida, J., et al., Mimicry of CD40 signals by Epstein-Barr virus LMP1 in B lymphocyte responses. Science, 1999. 286(5438): p. 300–3.

34. Shair, K.H., et al., EBV latent membrane protein 1 activates Akt, NFkappaB, and Stat3 in B cell lymphomas. PLoS Pathog, 2007. 3(11): p. e166.

35. Mainou, B.A., D.N. Everly, Jr., and N. Raab-Traub, Unique signaling properties of CTAR1 in LMP1-mediated transformation. J Virol, 2007. 81(18): p. 9680–92.

36. Mainou, B.A., D.N. Everly, Jr., and N. Raab-Traub, Epstein-Barr virus latent membrane protein 1 CTAR1 mediates rodent and human fibroblast transformation through activation of PI3K. Oncogene, 2005. 24(46): p. 6917–24.

37. Cahir-McFarland, E.D., et al., Role of NF-kappa B in cell survival and transcription of latent membrane protein 1-expressing or Epstein-Barr virus latency III-infected cells. J Virol, 2004. 78(8): p. 4108–19.

38. Luftig, M., et al., Epstein-Barr virus latent infection membrane protein 1 TRAF-binding site induces NIK/IKK alpha-dependent noncanonical NF-kappaB activation. Proc Natl Acad Sci U S A, 2004. 101(1): p. 141–6.

39. Sylla, B.S., et al., Epstein-Barr virus-transforming protein latent infection membrane protein 1 activates transcription factor NF-kappaB through a pathway that includes the NF-kappaB-inducing kinase and the IkappaB kinases IKKalpha and IKKbeta. Proc Natl Acad Sci U S A, 1998. 95(17): p. 10106–11.

40. Giehler, F., et al., Epstein-Barr virus-driven B cell lymphoma mediated by a direct LMP1-TRAF6 complex. Nat Commun, 2024. 15(1): p. 414.

41. Greenfeld, H., et al., TRAF1 Coordinates Polyubiquitin Signaling to Enhance Epstein-Barr Virus LMP1-Mediated Growth and Survival Pathway Activation. PLoS Pathog, 2015. 11(5): p. e1004890.

42. Izumi, K.M., et al., The Epstein-Barr virus oncoprotein latent membrane protein 1 engages the tumor necrosis factor receptor-associated proteins TRADD and receptor-interacting protein (RIP) but does not induce apoptosis or require RIP for NF-kappaB activation. Mol Cell Biol, 1999. 19(8): p. 5759–67.

43. Izumi, K.M. and E.D. Kieff, The Epstein-Barr virus oncogene product latent membrane protein 1 engages the tumor necrosis factor receptor-associated death domain protein to mediate B lymphocyte growth transformation and activate NF-kappaB. Proc Natl Acad Sci U S A, 1997. 94(23): p. 12592–7.

44. Izumi, K.M., K.M. Kaye, and E.D. Kieff, The Epstein-Barr virus LMP1 amino acid sequence that engages tumor necrosis factor receptor associated factors is critical for primary B lymphocyte growth transformation. Proc Natl Acad Sci U S A, 1997. 94(4): p. 1447–52.

45. Meckes, D.G., Jr., N.F. Menaker, and N. Raab-Traub, Epstein-Barr virus LMP1 modulates lipid raft microdomains and the vimentin cytoskeleton for signal transduction and transformation. J Virol, 2013. 87(3): p. 1301–11.

46. Katano, H., L. Pesnicak, and J.I. Cohen, Simvastatin induces apoptosis of Epstein-Barr virus (EBV)-transformed lymphoblastoid cell lines and delays development of EBV lymphomas. Proc Natl Acad Sci U S A, 2004. 101(14): p. 4960–5.

47. Lam, N. and B. Sugden, LMP1, a viral relative of the TNF receptor family, signals principally from intracellular compartments. EMBO J, 2003. 22(12): p. 3027–38.

48. Ardila-Osorio, H., et al., TRAF interactions with raft-like buoyant complexes, better than TRAF rates of degradation, differentiate signaling by CD40 and EBV latent membrane protein 1. Int J Cancer, 2005. 113(2): p. 267–75.

49. Gewurz, B.E., et al., Canonical NF-kappaB activation is essential for Epstein-Barr virus latent membrane protein 1 TES2/CTAR2 gene regulation. J Virol, 2011. 85(13): p. 6764–73.

50. DeKroon, R.M., et al., Global Proteomic Changes Induced by the Epstein-Barr Virus Oncoproteins Latent Membrane Protein 1 and 2A. mBio, 2018. 9(3).

51. Shair, K.H. and N. Raab-Traub, Transcriptome changes induced by Epstein-Barr virus LMP1 and LMP2A in transgenic lymphocytes and lymphoma. mBio, 2012. 3(5).

52. Minamitani, T., et al., Mouse model of Epstein-Barr virus LMP1- and LMP2A-driven germinal center B-cell lymphoproliferative disease. Proc Natl Acad Sci U S A, 2017. 114(18): p. 4751–4756.

53. Jiang, C., et al., CRISPR/Cas9 Screens Reveal Multiple Layers of B cell CD40 Regulation. Cell Rep, 2019. 28(5): p. 1307–1322 e8.

54. Liao, Y., et al., Lysine-specific histone demethylase complex restricts Epstein-Barr virus lytic reactivation. Nat Microbiol, 2025. 10(12): p. 3290–3304.

55. Sanson, K.R., et al., Optimized libraries for CRISPR-Cas9 genetic screens with multiple modalities. Nat Commun, 2018. 9(1): p. 5416.

56. Devergne, O., et al., Role of the TRAF binding site and NF-kappaB activation in Epstein-Barr virus latent membrane protein 1-induced cell gene expression. J Virol, 1998. 72(10): p. 7900–8.

57. Gewurz, B.E., et al., Genome-wide siRNA screen for mediators of NF-kappaB activation. Proc Natl Acad Sci U S A, 2012. 109(7): p. 2467–72.

58. Le Clorennec, C., et al., Molecular basis of cytotoxicity of Epstein-Barr virus (EBV) latent membrane protein 1 (LMP1) in EBV latency III B cells: LMP1 induces type II ligand-independent autoactivation of CD95/Fas with caspase 8-mediated apoptosis. J Virol, 2008. 82(13): p. 6721–33.

59. Choi, I.K., et al., Signaling by the Epstein-Barr virus LMP1 protein induces potent cytotoxic CD4(+) and CD8(+) T cell responses. Proc Natl Acad Sci U S A, 2018. 115(4): p. E686–E695.

60. Doench, J.G., et al., Optimized sgRNA design to maximize activity and minimize off-target effects of CRISPR-Cas9. Nat Biotechnol, 2016. 34(2): p. 184–191.

61. Zhang, B.C., et al., STEEP mediates STING ER exit and activation of signaling. Nat Immunol, 2020. 21(8): p. 868–879.

62. Liu, H.P., C.C. Wu, and Y.S. Chang, PRA1 promotes the intracellular trafficking and NF-kappaB signaling of EBV latent membrane protein 1. EMBO J, 2006. 25(17): p. 4120–30.

63. Izumi, K.M., K.M. Kaye, and E.D. Kieff, Epstein-Barr virus recombinant molecular genetic analysis of the LMP1 amino-terminal cytoplasmic domain reveals a probable structural role, with no component essential for primary B-lymphocyte growth transformation. J Virol, 1994. 68(7): p. 4369–76.

64. Verweij, F.J., et al., Exosomal sorting of the viral oncoprotein LMP1 is restrained by TRAF2 association at signalling endosomes. J Extracell Vesicles, 2015. 4: p. 26334.

65. Mosialos, G., et al., The Epstein-Barr virus transforming protein LMP1 engages signaling proteins for the tumor necrosis factor receptor family. Cell, 1995. 80(3): p. 389–99.

66. Kaye, K.M., et al., An Epstein-Barr virus that expresses only the first 231 LMP1 amino acids efficiently initiates primary B-lymphocyte growth transformation. J Virol, 1999. 73(12): p. 10525–30.

67. Hurwitz, S.N., et al., CD63 Regulates Epstein-Barr Virus LMP1 Exosomal Packaging, Enhancement of Vesicle Production, and Noncanonical NF-kappaB Signaling. J Virol, 2017. 91(5).

68. Flanagan, J., J. Middeldorp, and T. Sculley, Localization of the Epstein-Barr virus protein LMP 1 to exosomes. J Gen Virol, 2003. 84(Pt 7): p. 1871–1879.

69. Figueroa, C., J. Taylor, and A.B. Vojtek, Prenylated Rab acceptor protein is a receptor for prenylated small GTPases. J Biol Chem, 2001. 276(30): p. 28219–25.

70. Hutt, D.M., et al., PRA1 inhibits the extraction of membrane-bound rab GTPase by GDI1. J Biol Chem, 2000. 275(24): p. 18511–9.

71. Ma, M., et al., TAK1 is an essential kinase for STING trafficking. Mol Cell, 2023. 83(21): p. 3885–3903 e5.

72. Shair, K.H., et al., Epstein-Barr virus-encoded latent membrane protein 1 (LMP1) and LMP2A function cooperatively to promote carcinoma development in a mouse carcinogenesis model. J Virol, 2012. 86(9): p. 5352–65.

73. Zhang, B., et al., Immune surveillance and therapy of lymphomas driven by Epstein-Barr virus protein LMP1 in a mouse model. Cell, 2012. 148(4): p. 739–51.

74. Dirmeier, U., et al., Latent membrane protein 1 of Epstein-Barr virus coordinately regulates proliferation with control of apoptosis. Oncogene, 2005. 24(10): p. 1711–7.

75. Kilger, E., et al., Epstein-Barr virus-mediated B-cell proliferation is dependent upon latent membrane protein 1, which simulates an activated CD40 receptor. EMBO J, 1998. 17(6): p. 1700–9.

76. Burton, E.M., et al., Epstein-Barr virus latent membrane protein 1 subverts IMPDH pathways to drive B-cell oncometabolism. PLoS Pathog, 2025. 21(5): p. e1013092.

77. Meerbrey, K.L., et al., The pINDUCER lentiviral toolkit for inducible RNA interference in vitro and in vivo. Proc Natl Acad Sci U S A, 2011. 108(9): p. 3665–70.

78. Shalem, O., et al., Genome-scale CRISPR-Cas9 knockout screening in human cells. Science, 2014. 343(6166): p. 84–87.

79. Butler, T.J. and S.M. Smith, Strategies for the Purification of Membrane Proteins. Methods Mol Biol, 2023. 2699: p. 477–491.

80. Sun, Y., et al., Epstein-barr virus latent membrane protein 1 targets cIAP1, cIAP2 and TRAF2 for proteasomal degradation to activate the non-canonical NF-kappaB pathway. PLoS Pathog, 2026. 22(1): p. e1013898.

