## Supplement figures for "A CRISPR-Cas9 screen Reveals STEEP1 as a Key Host Dependency Factor for Epstein-Barr Virus Latent Membrane Protein 1 Trafficking and Signaling"

### Figure S1

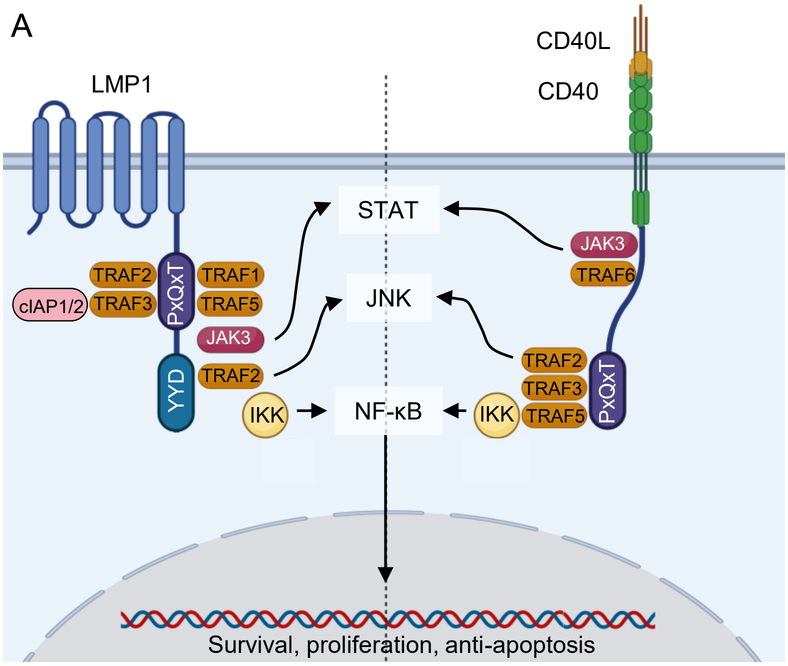
A
LMP1
CD40L
CD40
STAT
JNK
NF-κB
TRAF2
TRAF1
TRAF3
TRAF5
cIAP1/2
PxQxT
JAK3
YYD
TRAF2
IKK
JAK3
TRAF6
TRAF2
TRAF3
TRAF5
IKK
PxQxT
Survival, proliferation, anti-apoptosis

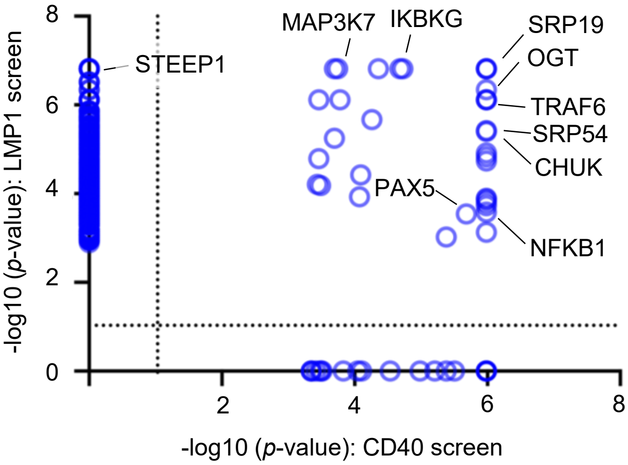
B
-log10 (p-value): LMP1 screen
-log10 (p-value): CD40 screen
0
2
4
6
8
STEEP1
MAP3K7
IKBKG
SRP19
OGT
TRAF6
SRP54
CHUK
PAX5
NFKB1

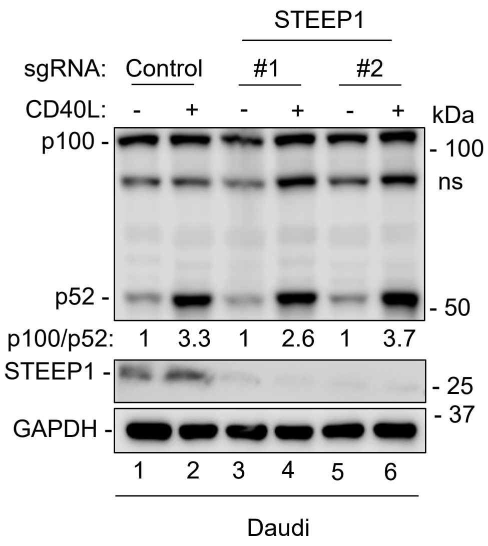
C
STEEP1
sgRNA: Control #1 #2
CD40L: - + - + - +
kDa
p100 -
- 100
ns
p52 -
- 50
p100/p52: 1 3.3 1 2.6 1 3.7
STEEP1 -
- 25
- 37
GAPDH -
1 2 3 4 5 6
Daudi

**Figure S1. STEEP1 is an LMP1 specific hit.**
(A) Schematic showing overlapping signaling pathways of LMP1 and CD40 in B cells.
(B) Scatter plot using the –log10(p-value) from the first and second replicate. The p-value cut off is 0.01.
(C) Immunoblot of Daudi B cells stimulated by CD40L with presence or absence of STEEP1. Daudi B cells were depleted STEEP1 and were stimulated by CD40L (50 ng/mL) for 48 hours. Blots represent n = 3 independent biological replicates.

### Figure S2

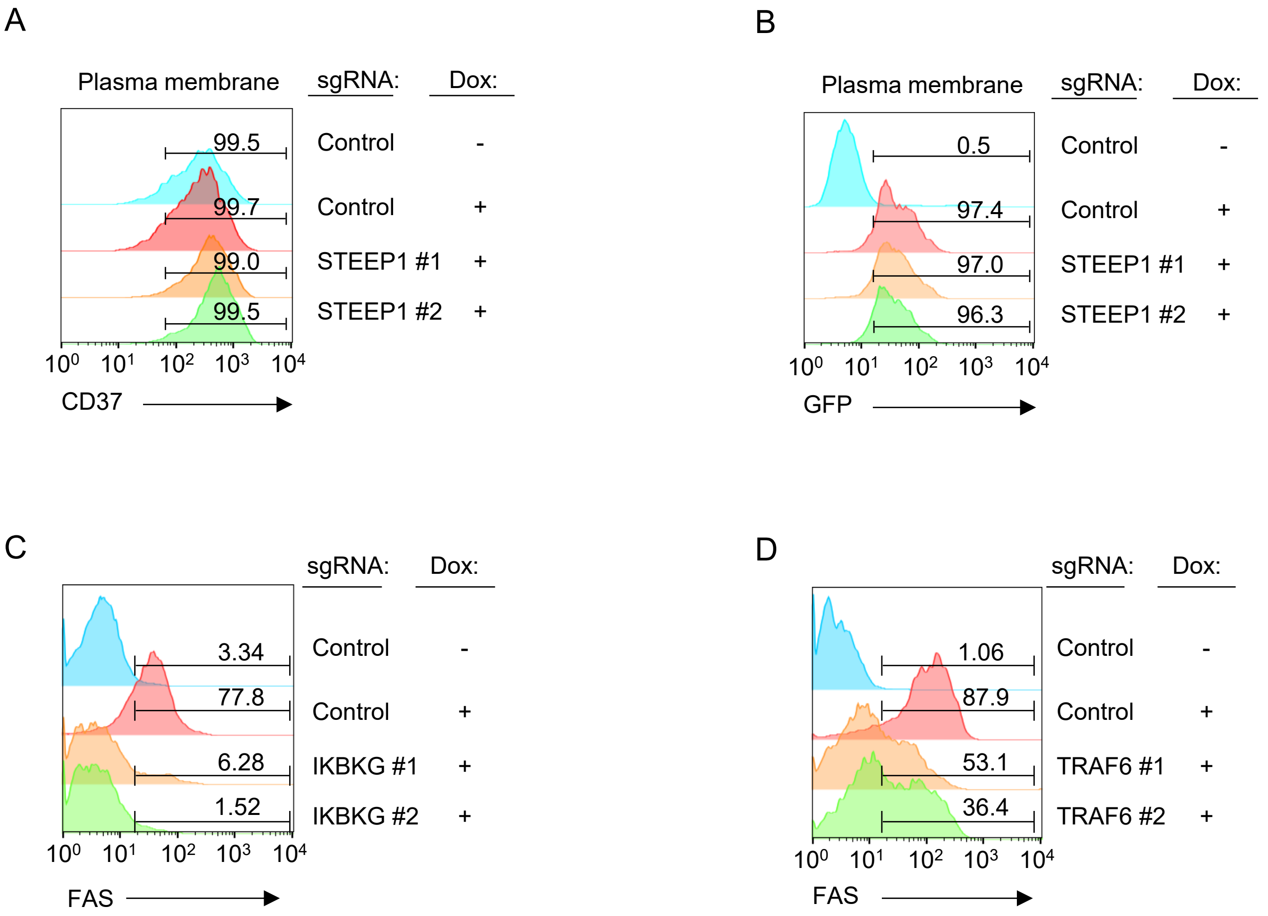
A
Plasma membrane
sgRNA:
Dox:
99.5
99.7
99.0
99.5
Control -
Control +
STEEP1 #1 +
STEEP1 #2 +
CD37
B
Plasma membrane
sgRNA:
Dox:
0.5
97.4
97.0
96.3
Control -
Control +
STEEP1 #1 +
STEEP1 #2 +
GFP
C
sgRNA:
Dox:
3.34
77.8
6.28
1.52
Control -
Control +
IKBKG #1 +
IKBKG #2 +
FAS
D
sgRNA:
Dox:
1.06
87.9
53.1
36.4
Control -
Control +
TRAF6 #1 +
TRAF6 #2 +
FAS

**Figure S2 STEEP1 depletion does not affect overall protein expression or membrane trafficking.**

(A) The plasma membrane expression of CD37, which is not an LMP1-target protein, was measured in Daudi cells expressing control single guide or two STEEP1 single guides. FACS chart is a representation of n = 3 biological replicates. The lines display the gate drawn. Numbers demonstrate the percentage of cells expressing plasma membrane CD37.

(B) The expression of GFP, which is engineered downstream of a Dox-responsible promoter, was measured in Daudi cells expressing control single guide or two STEEP1 single guides. FACS chart is a representation of n=3 biological replicates. The lines display the gate drawn. Numbers demonstrate the percentage of cells expressing GFP.

(C) The plasma membrane expression of Fas, which is upregulated by LMP1, was measured in Daudi cells expressing control single guide or two IKBKG single guides. IKBKG was one of the top screen hits. FACS chart is a representation of n = 3 biological replicates. The lines display the gate drawn. Numbers demonstrate the percentage of cells expressing plasma membrane Fas.

(D) The plasma membrane expression of Fas, which is upregulated by LMP1, was measured in Daudi cells expressing control single guide or two TRAF6 single guides. TRAF6 was one of the top screen hits. FACS chart is a representation of n = 3 biological replicates. The lines display the gate drawn. Numbers demonstrate the percentage of cells expressing plasma membrane Fas.

### Figure S3

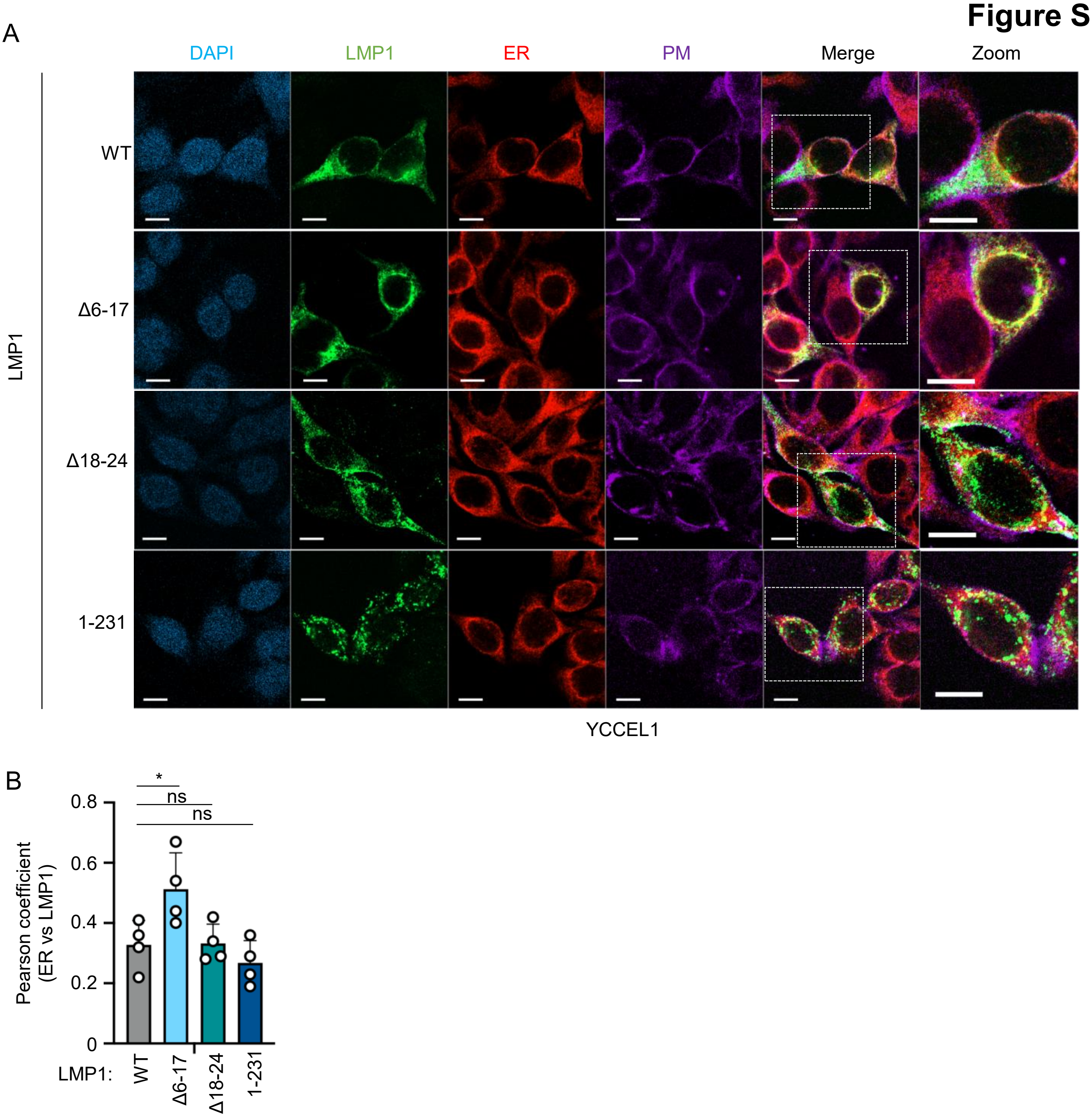
A
DAPI
LMP1
ER
PM
Merge
Zoom
LMP1
WT
Δ6-17
Δ18-24
1-231
YCCEL1
B
*
ns
ns
0.8
0.6
0.4
0.2
0
Pearson coefficient
(ER vs LMP1)
LMP1:
WT
Δ6-17
Δ18-24
1-231

**Figure S3 LMP1 N-terminal deletion affects LMP1 trafficking outside of ER.**

(A) Immunofluorescent microscopy images of YCCEL1 LMP1 WT and N-terminal domain mutants along with ER co-staining. YCCEL1 cells were transduced with a Dox-inducible LMP1 (WT or a various N-terminal truncation mutants). After puromycin selection for almost 2 weeks, cells were confirmed LMP1 expression by Western blot and seeded onto cover slips. The expression of LMP1 was turned on with Dox for 16 hours (200 ng/mL). Cells were then washed with PBS and stained for PM before fixation and permeabilization, and were stained for LMP1 (green) and ER (red). Scale bar = 10 μm.

(B) Colocalization analyses of ER and LMP1 (WT or N-terminal truncation mutants) in YCCEL1 cells. Images were analyzed by ImageJ using the colo2 plug-in. Pearson coefficient was calculated from each cell. *, statistically significant (One-way ANOVA, $p<0.05$), ns, non-significant.
